# Surfactant-mediated effects on hydrological and physical soil properties: a data synthesis

**DOI:** 10.1101/2023.07.02.547370

**Authors:** Anika Lehmann, Maximilian Flaig, Juan F. Dueñas, Matthias C. Rillig

## Abstract

Soils are under threat of a multitude of anthropogenic factors affecting the complex interplay of various physical and hydrological soil processes and properties. One such factor is the group of surface-active compounds. Surfactants have a broad range of applications, and can reduce solid-liquid interfacial forces and increase wettability and dispersion of particles. Surfactant effects are context-dependent, giving rise to a wide range of reported effects on different soil processes and properties.

Here, we evaluate the evidence base of surfactant research on 11 hydrological and physical soil variables. Our goal was to identify knowledge gaps and to test the robustness of proposed surfactant effects.

We found that the current knowledge base is insufficient to reach strong data-backed conclusions about effects of surfactants in soils. We identified a unique case of bias in the data as a result of conflated patterns of lab and field studies. We could not support the hypothesis that surfactant charge determines soil effects for any of the tested soil variables.

We believe that further experiments on surfactant mediated effects on soil properties and processes are urgently required, paying attention in particular to improving experimental design and data reporting standards.

**Graphical abstract:** 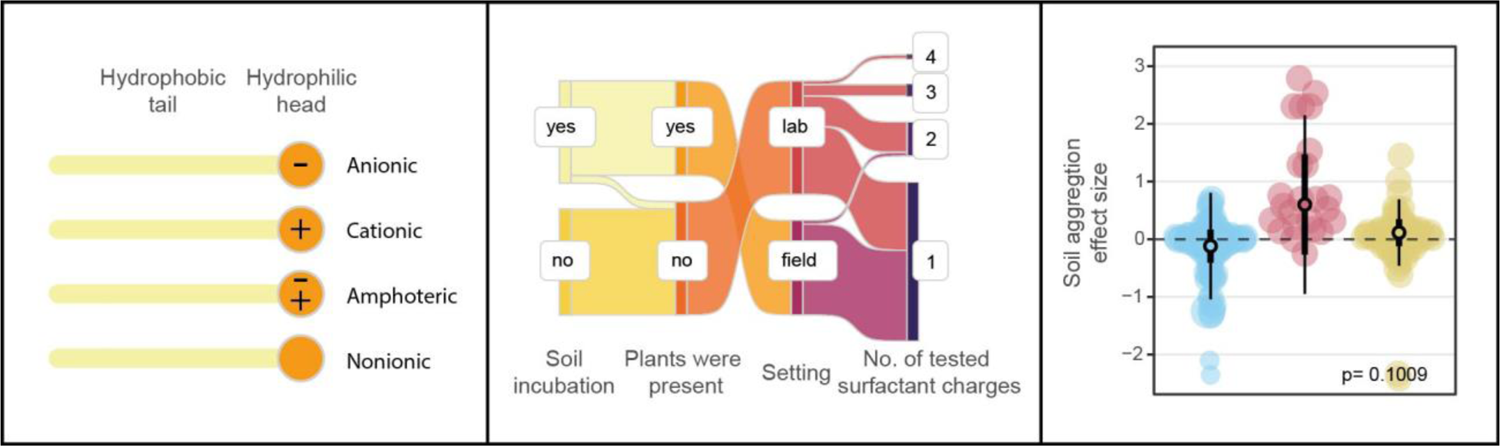

## 1. Introduction

Maintaining soil attributes and functions is crucial for sustaining soil health and the management of ecosystems (Lehmann *et al*., 2020). A key soil property is soil structure, the three-dimensional arrangement of soil particles and the resulting pore space (Six *et al*., 2004; Hao *et al*., 2008). Soil aggregation, the process resulting in soil structure, is affected by biological, chemical and physical factors and forces of natural and anthropogenic origin influencing component processes of soil aggregation (formation, stabilization, disintegration). As a result of changes in soil structure, various interlinked physical and hydrological soil processes and properties can be affected. For example, enrichment of soil with hydrophobic organic compounds can increase soil hydrophobicity, stabilize soil aggregates while reducing soil water infiltration capacity and increasing water run-off on the soil surface leading to soil erosion (Doerr *et al*., 2000; Shakesby *et al*., 2000). Soil erosion can cause aggregate breakdown and clogging of soil pores by fine particles, decreasing hydraulic conductivity and water infiltration to deeper soil layers. Soil pore clogging can restrict soil capillarity causing upper soil layers to dry more quickly, which may result in increased soil hydrophobicity (Kuhnt, 1993). Several synthetic organic chemical pollutants, and other global change factors, have the potential to affect this complex interplay of soil structure and other soil processes and properties, and one such group are surfactants.

Surfactants are surface active agents with a polar (hydrophilic) and apolar (hydrophobic) moiety, rendering them amphiphilic. At specific concentrations (critical micelle concentration) (Fuguet *et al*., 2005), surfactant monomers start to form micelles. The hydrophobic moiety points towards the micelle core while the hydrophilic moiety is oriented towards the aqueous interface. Their amphiphilic character and this subsequent micellization process make surfactants potent detergents and solvents, for example for hydrophobic organic compounds (Haigh, 1996). Furthermore, surfactants can affect surface characteristics by adsorption, which can cause hydrophobic particles to be covered by surfactant monomers, forming a hydrophilic layer (or vice versa) (Law *et al*., 1966; Ishiguro & Koopal, 2016). Hence, surfactants can reduce solid-liquid interfacial forces and increase wettability and dispersion of particles (Koopal, 2012).

There are five major groups of surfactants characterized by the charge of their polar moiety as a result of negatively or positively charged or non-ionized functional groups: anionic, cationic, nonionic, amphoteric (zwitterionic) and semi-polar (Le Toumelin & Baboulene, 1996; Zhu *et al*., 2018; Badmus *et al*., 2021).

The electrical charge of the polar and the molecular structure of both the polar and apolar moieties are key features of surfactants and determine their field of application and functional efficiency. In addition to being used as medical adjuvants and detergents in every-day hygiene products, surfactants are also frequently applied as wetting, emulsifying, and dispersing agents in the textile and agricultural industries. In soil remediation activities they are used as soil washing reagents for crude oil, diesel and light non-aqueous phase liquids decontamination (Ji *et al*., 2021) and as soil conditioners (e.g. wetting agents) to improve soil physical properties (Letey, 1975). That is, they are used to reduce water repellency, water run-off and soil erosion in natural (e.g. forests; Rice & Grismer, 2010) and urban areas (e.g. turfs and golf courses; Barton & Colmer, 2011).

Due to their wide range of application, various sources of soil contamination by surfactants are possible. The most common entry points into the environment occur via gray- and wastewaters and use of agrochemicals. Surfactant concentrations in graywater effluents can range from 0.7 to 70 mg L-1 (Wiel-Shafran *et al*., 2006) and in sewage sludges from 0.2 to 20 g kg-1, causing soil concentrations of up to several mg per kg soil (Kuhnt, 1993).

Surfactants are degradable in soil (Waters *et al*., 1989; Knaebel *et al*., 1990; Ying, 2006) but degrade more slowly than in aquatic environments (Kuhnt, 1993). They degrade faster under aerobic than under anaerobic conditions (Swisher, 1970). The chemical structure of the surfactant and physicochemical soil parameters (Cowan-Ellsberry *et al*., 2014; Arora *et al*., 2022) further modulate their biodegradability. The breakdown products of surfactants can be more persistent and toxic than the original surfactant product (Shang *et al*., 1999; Scott & Jones, 2000; Li *et al*., 2017; Badmus *et al*., 2021).

The toxicity of surfactants (and their breakdown products) depends on the chemical structure, the investigated target organisms (e.g. plants, bacteria and fungi;Gloxhuber, 1974; Rebello *et al*., 2014; Arora *et al*., 2022), and the environmental factors affecting the adsorption and hence mobility of the surfactant (Jahan *et al*., 2008).

In addition to their potential toxicity, it is consensus that surfactants alter soil characteristics (Kuhnt, 1993; Arora *et al*., 2022); for instance, repeated application of nonionic surfactants can cause soils to become water repellent (Song *et al*., 2021) because the surfactants form hydrophobic surfaces on soil particles.

The reported effects of surfactants on soil processes and properties are context dependent (e.g. for soil aggregation see Sutherland & Ziegler, 1998) with several key parameters influencing the surfactant-mediated effects on soil: (1) The charge of the polar moiety determines how well a surfactant adsorbs to soils (e.g. clay, organic matter and other negatively charged particles). Anionic surfactants adsorb less to negatively charged particles and compounds than cationic surfactants (Law & Kunze, 1966). Depending on the surfactant charge, its concentration and the sorptive capacity of the soil, more cationic and to a lesser degree nonionic surfactants can become adsorbed, which can lead to more hydrophobic surfaces in soil. While anionic surfactants remain mainly in the aqueous phase, they can reduce the surface tension of water and thus increase the wettability of soil (Kuhnt, 1993). (2) The soil chemistry and hence the soil texture, the amount and type of clay (Law & Kunze, 1966), pH (Xu *et al*., 1991) and organic matter content influence adsorption mechanisms in soils. For example, anionic surfactants can be adsorbed to a limited extent on kaolinite but not montmorillonite with adsorption being further reduced under higher soil pH (Xu *et al*., 1991). (3) Microbes can degrade surfactants and thus modulate their persistence and efficiency, while surfactants in turn affect the microbial community composition (Mandic *et al*., 2006; Olkowska *et al*., 2014) which can have ripple-on effects on soil functions and processes.

This context dependency leads to diverse research outcomes, so far limiting our ability to generalize (Kuhnt, 1993). Our goal here is therefore to evaluate the evidence base of surfactant research on hydrological and physical soil properties and processes, to identify any knowledge gaps, and to challenge the robustness of proposed surfactant effects.

We here focus on 11 variables determining the functionality of soil: Soil aggregation, soil porosity, clay dispersion, water repellency, water run-off, soil loss, water infiltration, water retention, water content, hydraulic conductivity and capillary rise.

Our approach is to search the literature for experiments on surfactant application affecting these 11 soil variables and to build a database to (1) investigate the reporting on key surfactant, soil and test system parameters by systematic mapping and (2) to evaluate surfactant-mediated effects reported in the literature by meta-analysis.

## 2. Methods

### 2.1. Search string development

We searched the Web of Science (WoS) Core Collection with default settings. We built a preliminary search string (supplementary materials, May 2020 with 328 hits) to collect a first set of potential matching articles for this project. The bibliometric data of these articles was exported to biblioshiny from the bibliometrix package in R (Aria & Cuccurullo, 2017) for an analysis of the most frequent words in titles, abstracts, author keywords and keywords plus to identify further useful terms. The final search string had different modules: the surfactant terms and synonyms, terms to filter for soil systems, terms representing soil properties of interest defined in the introduction, exclusion terms to reduce the number of mismatching articles (e.g. excluding biosurfactants) and WoS research areas to be excluded (mismatching WoS research areas were known from previous work from our lab). The final search string combined the 5 modules in a topic search:

TS = ((“surfactant*” OR “surface active agent*” OR “synthetic* detergent*” OR “soil conditioner$”) AND (“soil” OR “soils” OR “terrestrial”) AND ((“wetting” OR “wettable” OR “repellency” OR “hydraulic conductivity” OR “infiltration” OR “aggregat*” OR “erodibility” OR “hydrophobicity” OR “soil loss” OR “soil water content” OR “stability” OR “erosion” OR “runoff” OR “run-off” OR “porosity” OR “capillary rise”) NEAR/5 (“soil” OR “soils” OR “sand” OR “sands” OR “clay” OR “clays”)) NOT (*exclusion terms*) NOT (*WoS research areas*))

The complete search string is available in the supplementary materials. Only 3 articles of our first set of potential matching articles could not be retrieved by this search string because there were no abstracts available in WoS for these articles; in general, the lack of an abstract in the WoS database makes the capture of articles challenging. The final search was conducted on 18.02.2023 and yielded 346 hits. These articles were exported and screened for matching our inclusion/exclusion criteria. Studies had to present data on experiments with a surfactant treatment and corresponding surfactant-free control in a soil system (no aquatic studies, no pure clay or pure sand as substrate). If the surfactant charge was not reported or could not be obtained, then this study from an article was excluded. We also excluded all experiments on soil conditioners and wetting agents that were not surfactants (Zontek & Kostka, 2012), and also studies focusing on soil washing, sorption, co-contamination and transmission of co-contaminants through soil. Greywater and waste water treatments were not included, because no clear compound or product but rather a mixture of unknown surfactant and chemical composition and concentration is applied in these cases. Finally, experiments on biosurfactants (Mulligan, 2005; Bustamante *et al*., 2012) and soil amendments (e.g. manure, fertilizer) treatments were also excluded.

The screening resulted in 39 articles for which we screened the reference and citation list for further matches. We found additional 15 articles, among them the 3 articles of our first set of potential matching articles that could not be retrieved by the final search string. The 54 articles were used to build the database for the following analyses (Fig. S1). We followed the Prisma EcoEvo guideline for reporting standards (O’Dea et al., 2021).

### 2.2. Screening and coding

We included author name and publication year from the exported bibliometric data. Furthermore, we collected data on surfactant characteristics (name, charge, application concentration, critical micelle concentration, solubility in water), soil parameters (texture, clay content, organic matter content, pH, cation exchange capacity, water repellency of soil before the start of the experiment), experimental features (incubation time, presence of plants, setting (lab vs. field)) and information about soil properties of interest (treatment and control mean data, treatment and control variance data, sample size).

### 2.3. Systematic mapping

Based on the constructed data table, we built a summary table to investigate and visualize data availability and potential restrictions of the data set. Therefore, the data from each article were either coded as present/ absent or as counts (frequency of occurrence for tested surfactant charges and concentrations and soil types) per article. Only one article provided two studies (Fullen et al., 1993) reporting on results of a lab and a field experiment) and hence contributed two data rows to the systematic mapping analyses. All other articles contributed only one data row to the summary data table (see Fig.1 and 2; the summary data table is not the meta-analysis data table).

**Fig. 1.**
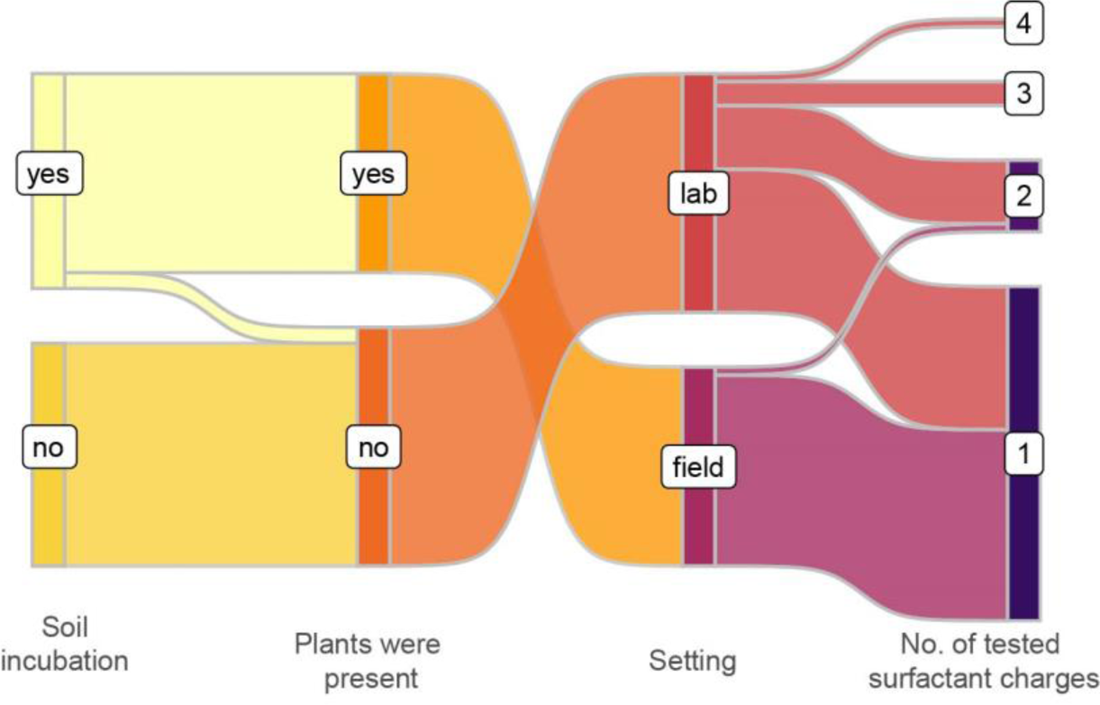
Sankey plot of experimental parameters of the 54 studies included (one study provided 2 experiments to this analysis resulting in 55 data rows). The four stages represent the key parameters of included experimental studies: 1) If the soil was incubated between treatment with surfactants and harvest and measurements, respectively; 2) If plants were present in the test system; 3) if studies were done in the field or under controlled conditions in the lab; 4) How many different surfactant charge (anionic, nonionic, cationic or zwitterionic) were tested in an experimental study. The nodes represent the frequency of occurrences for the different stage categories.

Figures were produced in R (version 4.2.2) with the packages ggplot2, ggpubr and ggsankey (Wickham, 2009; R Development Core Team, 2022; Sjoberg, 2023; Wickham, 2023).

### 2.4. Meta-analysis

Here we investigated the magnitude and variance of 11 hydrological/physical soil property and process effect sizes representing soil hydrological and structural response variables: soil aggregation, clay dispersion, soil porosity, (volumetric) water content, water retention, soil loss, wettability (water repellency), infiltration, capillary rise, hydraulic conductivity, water-run-off. The meta-analysis data table comprised all 54 articles which together provided 358 data rows.

For each of the 11 soil variables, we calculated log response ratio effect sizes (Yang *et al*., 2022) with the mean and variance data for treatment and control groups following the function: log response ratio = log(XT/XC). XT indicates the data of surfactant treated samples and XC the corresponding surfactant-free controls. We used the escalc() in the package “metafor” (Viechtbauer, 2010) to calculate the effect sizes and their variances by including the associated mean, standard deviation (sd) and sample size (*n*) data. If necessary, we converted standard error (se) data into sd data by the following function: sd= sqrt(n) x se. When data were presented in figures we used WebPlotDigitizer to obtain data (Rohatgi, 2022).

Several articles did not report either variance or sample size data (Fig. 2). In these cases, we applied data imputation techniques (Kambach *et al*., 2020). For each of the 11 soil variables, we applied specific techniques as they were necessary. For missing sample size data, we used the lowest detected sample size (n=3) from other articles. For missing variance data, we did median imputation if at least one article provided these data. If no variance data were available, we used the inverse of the sample size as variance surrogate (cases: clay dispersion and capillary rise).

**Fig. 2.**
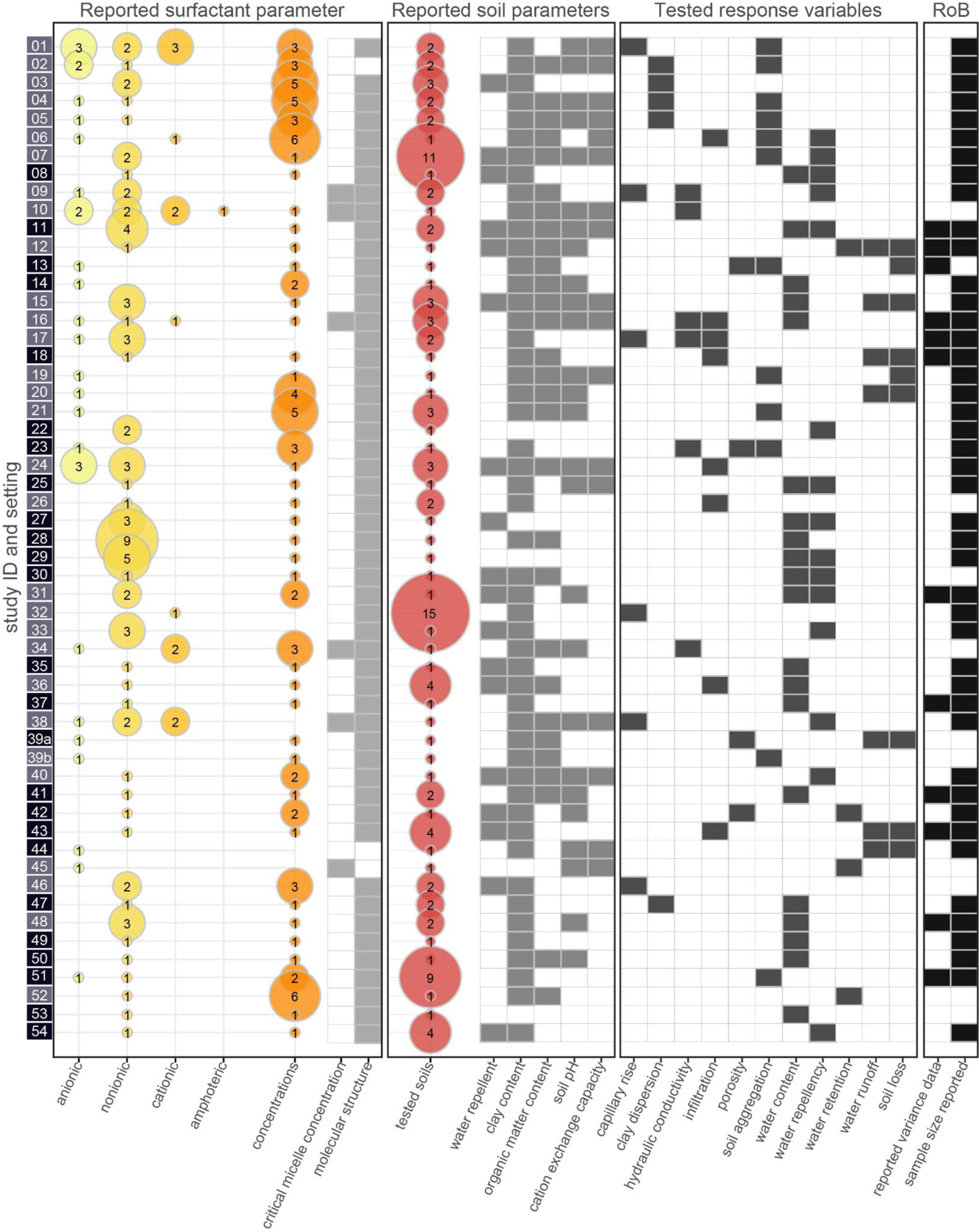
Balloon plot and binary heatmap for study parameters of the 54 studies included (one study provided 2 experiments to this analysis indicated with the letters a and b). The study identifier is underlaid with information on study setting (dark for field and light color for lab study). Reported surfactant parameters comprise surfactant charge (anionic, nonionic, cationic or zwitterionic), number of different concentrations tested and reporting of critical micelle concentration and molecular structure of tested surfactant(s). Reported soil parameters indicate the number of tested soils and reporting of key soil characteristics (if the soil was water repellent or not before the start of the treatment application, the clay and organic matter content, the soil pH and cation exchange capacity). Multiple studies tested one or multiple surfactant products on different soils. Tested response variables indicate which hydrological and physical soil variables are included in the data set. Risk of bias (RoB) reflects the reporting of critical statistical data (if variance and sample size data were reported or not). The balloons give frequency information for each category represented by their size; additionally, exact numbers are given as balloon overlays. The binary heatmaps indicate presence (filled tiles) or absence (white tiles) of specific data.

For the analysis, we focused on test system, surfactant, and soil characteristics with high data coverage per soil variable to limit restrictions due to low statistical power and biases. Setting was tested as categorical moderator with the two levels field and lab.

Surfactant charge was included as a categorical moderator with the three levels anionic, cationic and nonionic. The single case of a reported amphoteric (zwitterionic) surfactant was excluded from the meta-analysis.

Surfactant application concentration was a numerical moderator with the unit mg kg-1. We set 10,000 mg kg-1 as the upper concentration limit to be included in this analysis. The data base included an article with tested surfactant concentrations of 1 to 100,000 times the manufacturer-recommended rate (Ziegler & Sutherland, 1998). The surfactant concentration data were log10-transformed for improved data distribution.

Soil clay content was tested as numerical moderator with the unit %. In cases where no clay content but the soil texture following the USDA nomenclature was reported, we imputed mean clay content data derived from the USDA soil triangle.

For the statistics, we used mixed-effects meta-analysis with author cluster and study identifier as random factors to account for non-independency of multiple effect sizes per study and multiple studies published by the same author cluster. For the database, there was a cluster consisting of 3 papers and 11 clusters involving 2 papers with overlapping author/s.

We used the rma.mv() function in the “metafor” package with restricted maximum likelihood estimator and applied a t-distribution with k - p degrees of freedom, thereby mimicking the Knapp and Hartung method (Knapp & Hartung, 2003) for increased robustness of model outcomes against false positive results (Assink & Wibbelink, 2016); k and p are the number of effect size estimates and model coefficients being included in the model. For models with moderator variables, we fitted a heterogeneous variance model and set the covariance between moderator levels to zero (Viechtbauer, 2010). We implemented the Nelder-Mead method of optimization if models failed to converge (O’Dea *et al*., 2019).

For sensitivity analyses, we tested for publication bias via small-study effect and time-lag effect (Sterne *et al*., 2000; Sterne & Egger, 2001; Koricheva *et al*., 2013; Nakagawa *et al*., 2022). We included year of publication and standard error in a multi-moderator mixed-effects meta-analysis model (Nakagawa *et al*., 2021). A significant slope indicates a case of bias. Plots were produced with the “orchard 2.0” package (Nakagawa *et al*., 2020).

## 3. Results

### 3.1. Systematic mapping

We found a unique bias in the dataset (Fig. 1), resulting from the conflated pattern of study parameters of experiments in the field and the lab. In all field studies, plants were present and there was an incubation time between surfactant application and sample collection. For lab studies, plants were never a component of the test system and only in one study were the surfactant treated samples incubated (Fullen *et al*., 1993). Hence, any detected effects for the moderator ‘setting’ cannot be solely assigned to higher degree of environmental or experimental control in lab studies compared to field studies, but could also be a result of plant presence or absence and soil biota (e.g. microbes) activity during incubation time.

We also found that tests with multiple surfactant products and surfactants of different charges were performed primarily in lab studies (Fig.1), while in field studies nonionic surfactants were preferentially tested (20 of 25 field studies).

In general, study focus was either on the surfactant products, the impact of different concentrations or the modulating effect of different soils on the surfactant-mediated effects on the soil variables. There are studies testing up to 9 different surfactants for one concentration in one soil (Xiang *et al*., 2021), one surfactant with 6 different concentrations in one soil (Karagunduz *et al*., 2001; Qi *et al*., 2017) or one surfactant with one concentration in up to 15 different soils (Nagy & Deák, 2013).

The data quality was variable across the studies and the 11 variables (Fig. 2). The most articles could be retrieved for soil water content (20 articles) and water repellency (15 articles). Soil porosity and soil water retention were both reported in 4 articles. A maximum of 3 of our targeted 11 variables were tested together in one study. Tests of relationships among the 11 variables in this dataset via multivariate meta-analysis models is thus not possible due to lack of data.

The targeted surfactant and soil parameters were not reported for all studies. The surfactants were either labeled by their chemical or brand name. Many surfactant products are blends of different surfactants and other additives. An in depth analysis based on the chemical or molecular structure of the different products was not possible. The critical micelle concentration was reported in 6 of the 54 articles.

Among the soil parameters, data on clay content could be retrieved for 49 of 54 articles which made this parameter an interesting candidate for the meta-analysis. For the other parameters, the data availability was insufficient: soil organic matter was reported in 28 of 54 articles, soil pH in 26 of 54 articles, soil CEC in 17 of 54 articles and soil water repellency before the start of treatment in 18 of 54 articles.

The evaluation of sample size and variance data, specifically necessary for performing meta-analyses, revealed that variance data was reported in 12 of 54 articles. This affected all 11 variables. Sample size was missing in 10 of 54 articles. The highest sample sizes we detected ranged between 6 and 8, while overall the dataset was dominated by reported experiments with 3 (17 of 54 articles) or 4 (17 of 54 articles) replicates.

### 3.2. Meta-analysis

Following the identified specifics of the dataset, we decided to focus on the overall effect of surfactant application on the 11 hydrological and physical variables. Furthermore, we tested the influence of the setting, surfactant charge, surfactant concentration and soil clay content as these were the variables with most consistent reporting across all studies and all 11 soil variables.

Overall, the application of surfactant had no clear effect on 10 of the 11 variables. Only for the soil loss effect size we identified a negative effect (Fig. 3; point estimate = −0.5263; confidence interval = [−0.8083 to −0.2442]; prediction interval = [-1.1536 to 0.1010]; p-value = 0.0008); soil loss is reduced in surfactant treated samples as compared to untreated controls.

**Fig. 3.**
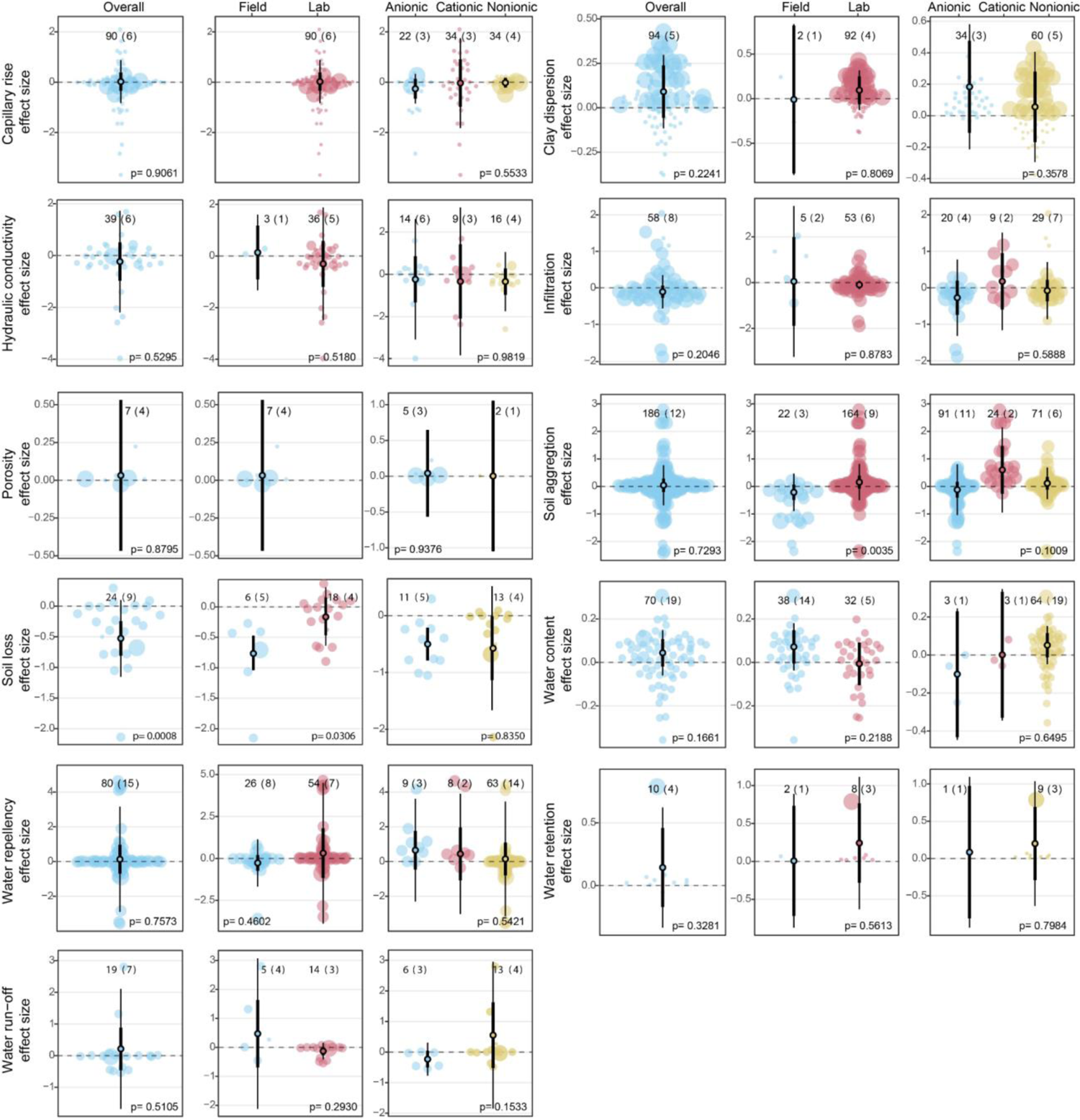
Overall, study setting and surfactant charge effects on individual soil hydrological and physical property and process effect sizes (capillary rise, clay dispersion, hydraulic conductivity, infiltration, porosity, soil aggregation, soil loss (splash and detachment erosion), water content, water repellency (hydrophobicity), water retention and water run-off; sorted by alphabet). Effect sizes and their variances are displayed as point estimate, 95% confidence (thick, black line) and prediction interval (thin, black line) with individual effect sizes estimates in the background. The number of effect sizes (k) included in each point estimate are given together with the numbers of articles (number in brackets) from which these k effect sizes originate. The dashed line represents the “line of no effect”. The p-value for the overall effect indicates if the effect is positive or negative (p-value < 0.05) or neutral (p-value > 0.05). For the moderator tests (setting and surfactant charge), the p-value indicates a statistical difference between the moderator levels (p-value > 0.05 represents no statistical difference). Model outcomes are given in Table S1 to S3.

For setting, we found that the point estimates for the soil aggregation and soil effect sizes are different for field and lab studies (Fig. 3; test of moderator p-value 0.0035 and 0.0306, respectively); the effect sizes are generally lower for field samples than for samples from laboratory experiments. A negative effect could only be detected for infiltration rate for lab studies (Fig. 3; point estimate = −0.09637; confidence interval = [-0.1745 to −0.0182], prediction interval = [-0.2861 to 0.0933]) and soil loss for field studies (Fig. 3; point estimate = −0.7559; confidence interval = [-1.0394 to −0.4724]; prediction interval = [-1.0394 to −0.47246]); surfactant treated soils showed faster water infiltration in the lab and reduced soil loss in the field compared to their respective controls.

Testing the surfactant charges revealed no clear differences among anionic, cationic and nonionic surfactants for all 11 evaluated hydrological and physical variables (Fig. 3). The general negative effects of surfactants on the soil loss effect size also manifested for the surfactant charges.

The surfactant concentration showed clear negative and positive effects for 6 of the 11 variables (Fig.S2; Table S4). We found negative relationships for soil aggregation and anionic surfactants, capillary rise for both anionic and cationic surfactants, water infiltration and anionic surfactants, hydraulic conductivity for both anionic and cationic surfactants, soil repellency for both cationic and nonionic surfactants and water run-off and anionic surfactants. Positive relationships were detected for soil aggregation and infiltration for both cationic and nonionic surfactants. The analyses were limited by the low number of articles testing surfactant concentrations. For some analysis data were derived from only one article; thus the results are not generalizable.

Analyzing the clay content of the test soil and its effect on the 11 variables showed negative and positive effects (Fig.S3; Table S5). Soil aggregation was reduced with increasing clay content when anionic and nonionic surfactants were applied compared to surfactant free controls. For hydraulic conductivity and anionic surfactants, as well as water repellency and cationic surfactants, we also found a negative effect with increasing clay content. For cationic surfactants, we found positive relationships between clay content and capillary rise, water infiltration and hydraulic conductivity effect sizes. For nonionic surfactants, this positive relationship was detected for hydraulic conductivity, water repellency and water run-off effect sizes.

It is worth noting that the majority of slopes showing a clear directional effect are close to zero indicating a limited biological relevance. The only exception was found for water repellency and cationic surfactant but this result is derived from data of just two articles providing 8 individual effect sizes (data points in the analysis). In general, the data for the clay content analysis are restricted due to limitations in the number of articles covering this topic.

We tested for publication biases. The publication year showed a positive relationship with the capillary rise effect size indicating that newer studies reported more positive results than older publications (Fig.S4; Table S6). For the soil aggregation effect size, a trend (p-value = 0.0584) for a negative relationship with the publication year was found, indicating that newer articles tend to report more negative results of surfactants on soil aggregation than older studies.

We detected small-study biases for the soil aggregation, infiltration and water repellency effect sizes (Fig.S5; Table S6). Studies with low sample size tend to report more negative results for soil aggregation and water repellency and more positive results for infiltration than studies with higher sample size and hence higher precision.

The analysis for time-lag bias furthermore revealed gaps along the publication timeline for multiple variables. Data on soil aggregation, soil loss, hydraulic conductivity and water repellency were published throughout the decades, while for the other variables no data were published between 1980 to 2000. Clay dispersion is a special case because after the year 2000 no more data were published for this variable.

## 4. Discussion

### 4.1. Systematic mapping

The systematic mapping of the literature on surfactant-mediated effects on the 11 soil variables revealed large knowledge gaps. Reporting standards varied among articles, and studies collectively addressed a limited parameter space.

Overall, it is surprising that our knowledge of soil effects for such a very commonly used and environmentally relevant group of synthetic organic chemicals is so limited. This likely reflects the general attitude that perhaps these effects are not important to study, maybe because these substances are widely regarded as readily biologically degradable.

The lack of incubation phases in lab studies, thus effectively ignoring microbial activity on surfactant-mediated effects, hampers realistic inferences on surfactant mediated effects on soil functionality and biodiversity (Fig. 1). Microbes can degrade surfactants and thus can promote production of breakdown products (Waters *et al*., 1989; Knaebel *et al*., 1990; Ying, 2006), but in turn microbes are also affected by surfactants and their breakdown products (Mandic *et al*., 2006; Olkowska *et al*., 2014). Neglecting these important processes rather drastically restricts our mechanistic understanding and thus any risk assessment of surfactants in soil.

Furthermore, there is also a lack of plants in lab studies. Plants can be affected by surfactants and their breakdown products (Cisar *et al*., 2000; Gálvez *et al*., 2019) and influence microbial activity, community composition and soil functions and processes (Bais *et al*., 2006).

Lab experiments are crucial for unraveling mechanisms underpinning effects of pollutants or environmentally relevant chemicals, because they provide the necessary levels of experimental control. It is thus of concern that lab experiments on this research topic do not fully cover some very important players, especially plants and microbes, necessary for properly evaluating surfactant-mediated effects. We thus urge that future studies explicitly include such aspects especially in lab experiments.

Context dependency is a challenge in studying surfactant-mediated effects, highlighting the necessity to represent different soil and other parameters, including the surfactants themselves. However, only a few studies tested a range of key parameters (e.g. soil and/or surfactant parameters) for multiple soil processes and properties. In the few cases where this was done, the studies were mainly conducted under lab conditions with the ‘setting bias’ discussed above. This raises the question to what extent lab study data can be generalized to field situations.

Surfactants have a wide range of chemical structures, likely giving rise to differences in interactions with abiotic and biotic factors in the test systems. Covering the whole parameter space is impossible, but focusing on key parameters allows concentrated research efforts to generate data for a more robust database (see Kuhnt, 1993). We strongly suggest improved reporting on the surfactant products and their chemical characteristics (e.g. including charge and critical micelle concentration), as well as soil parameters (e.g. clay content, organic matter content and soil hydrophobicity before surfactant application).

Furthermore, we note issues with overall data quality and reporting, in particular the reporting on essential statistical metrics like sample size and variance was limited (Fig. 2). We suggest that future studies pay attention to reporting these parameters to allow future data synthesis activities access to better data quality.

### 4.2. Meta-analysis

Due to the data quality and reporting issue highlighted above, the collated data allow only limited analyses of surfactant-mediated effects on soil processes and properties. Additionally, some of the 11 soil variables were addressed only by a limited number of articles, further restricting our overall capability to evaluate surfactant-mediated effects suggested in the literature. Thus we here focus on analyses with high statistical power.

We detected an overall negative impact of surfactants on the soil loss effect size, meaning that less soil is lost in surfactant treated samples during experimentally simulated erosion events (e.g. splash erosion by rain simulator) compared to the respective controls (Fig. 3). This overall effect was mainly driven by data from field studies and anionic surfactant application for which also a clear negative effect could be detected (Fig. 3). All studies focusing on the impact of anionic surfactants on soil loss used the product Agri-SC (e.g. Fullen et al., 1993) and hence the generalizability of this pattern is limited. As a soil conditioner, Agri-SC is suggested to reduce soil loss by improving water infiltration and soil crust formation (Lehrsch et al., 2011; Yönter & Yağmur, 2011).

We found an effect of the moderator ‘setting’ for the variables soil aggregation and soil loss. In field studies, the soil aggregation and soil loss effect sizes were lower compared to lab studies (Fig. 3). As summarized by Lehrsch et al. (2011), in lab studies unrealistically high treatment concentrations beyond cost-effective application rates for field production are tested. This could lead to a distorted picture for lab study data. Furthermore, as a consequence of the identified ‘setting bias’ (Fig.1) all field experiments experience an incubation phase compared to the majority of lab studies (Fullen et al., 1993) as the sole exception in this dataset). During an incubation phase, biological, chemical and physical soil factors are interacting and can be affected by the applied surfactant and its breakdown products to various degrees (Kuhnt, 1993). The lower effect sizes detected for soil aggregation and soil loss in field studies could be an indicator of the potential impact of an incubation phase and that we might overestimate surfactant-mediated effects on soil properties and processes by neglecting incubation phases in lab studies. This research gap needs to be closed in the future.

There was no detectable effect for surfactant charge, which is contradictory to claims in the literature. For example, for soil aggregation, we find a trend with anionic surfactants causing a decrease and nonionic surfactants an increase in the effect size values but this trend is not statistically supported (Table S3). In the literature, there are repeated reports and accepted hypotheses highlighting negative anionic and positive nonionic surfactant effect (e.g. Law *et al*., 1966; Mbagwu *et al*., 1993; Piccolo & Mbagwu, 1994): namely that anionic surfactants form hydrophilic, and nonionic surfactants hydrophobic, coatings around soil aggregates decreasing or increasing, respectively, aggregate protection against water infiltration and hence breakdown. Our results challenge the support for these established hypotheses. We clearly need more information on the context-dependence to arrive at such generalizations of surfactant-mediated effects on soil properties and functions.

An interesting summary of context-dependency for surfactant-mediated effects on soil hydraulic conductivity was reported by Tumeo (2007). This survey identified a total of 10 hypotheses which can support either an increase or a decrease in surfactant-mediated effects on soil hydraulic conductivity. Soils are complex, heterogeneous environments with their biotic (e.g. microbes, plants) and abiotic (e.g. clays) components interacting. Depending on the surfactant product and its properties the net effects can cover the whole range of positive, neutral and negative effects, which can then further differ spatially and temporarily. Surfactant-mediated effects are clearly context-dependent and the data available for this meta-analysis do not allow disentangling in which context what surfactant-mediated effect is to be expected.

We found effects for surfactant concentration and clay content on several of the 11 variables, but effects were either small and of low biological relevance or derived from a small number of data points of one or a few experiments. In some cases, studies tested high surfactant concentrations beyond expectable soil contamination levels (Kuhnt, 1993; Ziegler & Sutherland, 1998). Our cut-off threshold of 10,000 mg kg-1 is above recommended product concentrations. The identified relationships between surfactant concentrations and some soil variables might be exaggerating effects possible under realistic field conditions.

Nevertheless, in future studies, key characteristics like soil clay content and surfactant concentration should be considered to create a more solid database to allow meaningful analysis.

We detected some cases of bias: one case of time lag bias and three cases of small study bias. For capillary rise, newer studies tend to present more positive effect sizes than older studies. For soil aggregation and water repellency, studies with low numbers of replicates report lower effect sizes than studies with high sample size. The opposite is true for the infiltration effect size. These biases might be an artifact of the low number of articles for the majority of variables. But especially for soil aggregation there could be a special focus on anionic surfactants because of the well-established hypothesis on their negative effects on soil aggregates.

In conclusion, surfactants are present in many products and thus will be environmentally available in many soils. The current database is insufficient to reach strong data-backed conclusions about effects of surfactants in soils, and we believe that further testing is urgently required. We recommend that future research improve experimental design and data reporting. In particular, we urge researchers to report as much contextual information about tested soils as possible, as well as sample sizes and variance of results obtained. Finally, the indirect effects of surfactant presence on soil microbial organisms and plants need to be assessed.

## Acknowledgements.

We acknowledge support from the Einstein Research Unit “Climate and Water under Change” from the Einstein Foundation Berlin and Berlin University Alliance (grant no. ERU-2020-609). MCR acknowledges funding from an ERC Advanced Grant (694368).

## Author contributions

AL initiated the work. AL and MCR wrote the first draft. AL built the final dataset and conducted the systematic mapping. AL and MF conducted the meta-analysis. All authors (AL, MF, JFD and MCR) contributed to the writing of the manuscript.

