## supplementary material for "Surfactant-mediated effects on hydrological and physical soil properties: a data synthesis"

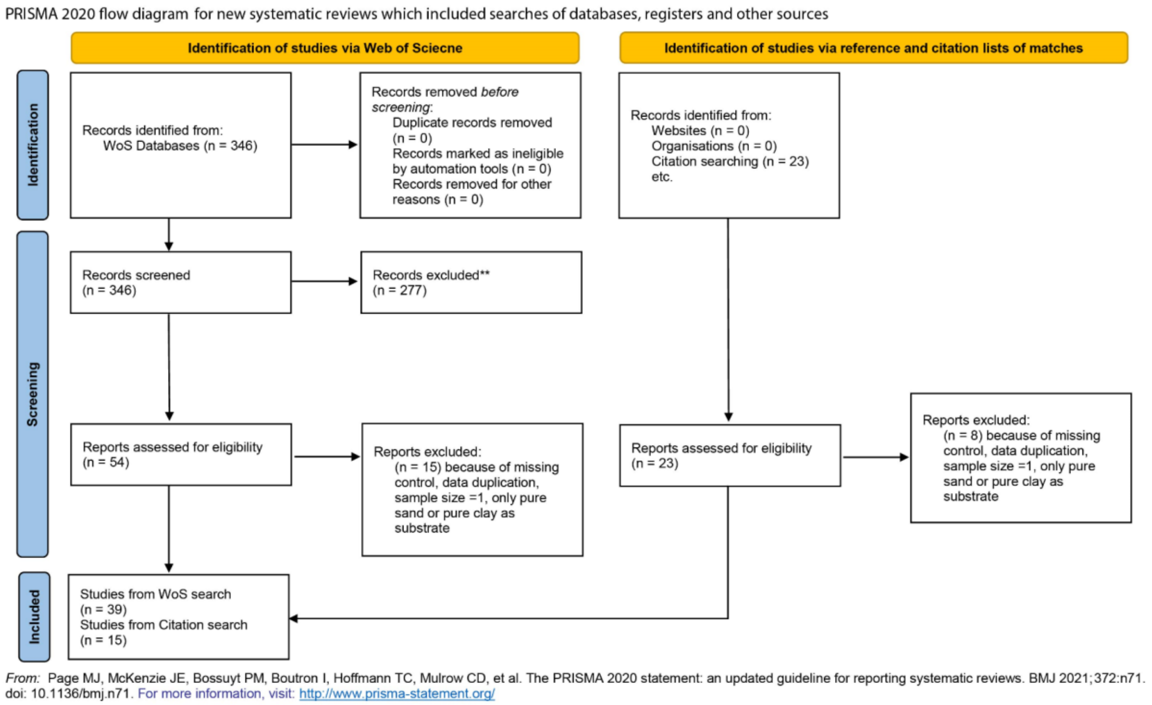

**Fig.S1.** PRISMA flow diagram

Preliminary search string

| search type | boolean operator | search string module | rationale |
|---|---|---|---|
| TS= | | ("Surfactant*" OR "surface active agent*" OR "surface active chemical*" OR "synthetic* detergent*" OR "soil conditioner$") | *surfactant terms and synonyms* |
| | AND | ("soil" OR "soils" OR "terrestrial") | *filter results towards soil systems* |
| | AND | ("runoff" OR "penetration*" OR "run-off" OR "wettability" OR "aggregate" OR stability" OR "hydraulic conductivity" OR "bulk density" OR "erodibility" OR "aggregate" NEAR/5 "stability") | *define potential soil properties as response variables of inetrest* |
| | NOT | ("compost" OR "vermicompost" OR "hydrogel" OR "biochar" OR "*litter" OR "*gypsum" OR "*manure" OR "contamination" OR "biopolymer" OR "biosurfactant" OR "biosurfactants" OR "bioemulsifier" OR "bioemulsifiers" OR "decontamination" OR "phytoremediation" OR "bioremediation" OR ("micelle*" NEAR/5 ("stability" OR "aggregation")) OR (("fluorosurfactant" OR "fluorosurfactants" OR "surfactant" OR "surfactants" OR "surface active") NEAR/100 ("sorption" OR "sorptive" OR "desorption" OR "adsorption" OR "adsorptive")) OR "drilling" OR "quenching" OR "biodegradation" OR "biodegraded" OR "copolymers" OR ((endogenous OR exogenous) NEAR/0 surfactant*) OR "lung" OR "lungs" OR "pulmonary" OR "alveolar" OR "nasal") | *define exclusion terms to reduce numbers of mismatching articles* |

Benchmark studies reference list

| First author last name | Publication title | Journal | Publication year |
|---|---|---|---|
| Qi | Effects of ionic surfactants on the aggregate stability and water repellency of silt loam soil | J soil sediments | 2017 |
| Lehrsch | Surfactant effects on the water-stable aggregation of wettable soils from the continental USA | hydrological processes | 2013 |
| Lehrsch | Surfactant effects on soil aggregate tensile strength | Geoderm | 2012 |
| Aamlid | The potential of a surfactant to restore turfgrass quality on a severely water-repellent golf green | Biologia | 2009 |
| Abu-Zreig | Effect of application of surfactants on hydraulic properties of soils | Biosystems Engineering | 2003 |
| Allred | SURFACTANT-INDUCED REDUCTIONS IN SOIL HYDRAULIC CONDUCTIVITY | Ground Water Monitoring and Remediation | 1994 |
| Barton | Granular wetting agents ameliorate water repellency in turfgrass of contrasting soil organic matter content | Plant and Soil | 2011 |
| Darboux | Evaluation of two soil conditioners for limiting post-fire erosion as part of a soil conservation strategy | Soil Use and Management | 2008 |
| Fullen | Effects of 'Agri-SC' soil conditioner on soil structure and erodibility: Some further observations | Soil Use and Management | 1995 |
| Kostka | Amelioration of water repellency in highly managed soils and the enhancement of turfgrass performance through the systematic application of surfactants | Journal of Hydrology | 2000 |
| Lehrsch | Surfactant and irrigation effects on wettable soils: runoff, erosion, and water retention responses | Hydrological Processes | 2011 |
| Mingorance | Laboratory methodology to approach soil water transport in the presence of surfactants | Colloids and Surfaces a-Physicochemical and Engineering Aspects | 2007 |
| Mobbs | Effects of four soil surfactants on four soil-water properties in sand and silt loam | Journal of Soil and Water Conservation | 2012 |
| Sullivan | Evaluating a nonionic surfactant as a tool to improve water availability in irrigated cotton | Hydrological Processes | 2009 |
| Sutherland | The influence of the soil conditioner 'Agri-SC' on splash detachment and aggregate stability | Soil & Tillage Research | 1998 |
| Yonter | Effect of Agri-SC as a soil conditioner on runoff, soil loss and crust strengths | African Journal of Biotechnology | 2011 |

| | | | |
|---|---|---|---|
| Ziegler | Effect of an anionic soil conditioner on water stable aggregation of three Hawaiian soils | Communications in Soil Science and Plant Analysis | 1998 |
| Cisar | The occurrence and alleviation by surfactants of soil-water repellency on sand-based turfgrass systems | Journal of Hydrology | 2000 |
| Barton | Ameliorating water repellency under turfgrass of contrasting soil organic matter content: Effect of wetting agent formulation and application frequency | | 2011 |
| Fitch | Effects of a Conditioner on Soil Physical Properties | Soil Science Society of America Journal | 1989 |
| Miyamoto | Effects of Wetting Agents on Water Infiltration into Poorly Wettable Sand, Dry Sod and Wettable Soils | Irrigation science | 1985 |
| Law | Reactions of Surfactants with Montmorillonitic Soils | SOIL SCI. SOC. AMEE. PHOC | 1966 |

Final search string

| Search type | Boolean operator | Search string module | Rationale |
|---|---|---|---|
| TS= | | (("surfactant*" OR "surface active agent*" OR "synthetic* detergent*" OR "soil conditioner$") | *surfactant terms and synonyms* |
| | AND | ("soil" OR "soils" OR "terrestrial") | *filter results towards soil systems* |
| | AND | (("wetting" OR "wettable" OR "repellency" OR "hydraulic conductivity" OR "infiltration" OR "aggregat*" OR "erodibility" OR "hydrophobicity" OR "soil loss" OR "soil water content" OR "stability" OR "erosion" OR "runoff" OR "run-off" OR "porosity" OR "capillary rise") NEAR/5 ("soil" OR "soils" OR "sand" OR "sands" OR "clay" OR "clays")) | *define potential soil properties as response variables of interest* |
| | NOT | ("biopolymer" OR "biosurfactant" OR "biosurfactants" OR "bioemulsifier" OR "bioemulsifiers" OR "decontamination" OR "phytoremediation" OR "bioremediation" OR ("micelles" NEAR/5 ("stability" OR "aggregation")) OR (("fluorosurfactant" OR "fluorosurfactants" OR "surfactant" OR "surfactants" OR "surface active") NEAR/100 ("sorption" OR "sorptive" OR "desorption" OR "adsorptive")) OR "drilling" OR "quenching" OR "biodegradation" OR "biodegraded" OR "copolymers" OR ((endogenous OR exogenous) NEAR/0 surfactant*) OR "lung" OR "lungs" OR "pulmonary" OR "alveolar" OR "nasal") ) | *define exclusion terms to reduce numbers of mismatching articles* |
| | NOT | SU=("Acoustics" OR "Allergy" OR "Anesthesiology" OR "Anthropology" OR "Archaeology" OR "Architecture" OR "Area Studies" OR "Art" OR "Arts & Humanities-Other Topics" OR "Asian Studies" OR "Audiology & Speech-Language Pathology" OR "Automation & Control Systems" OR "Biomedical Social Sciences" OR "Business & Economics" OR "Cardiovascular System & Cardiology" OR "Classics" OR "Communication" OR "Computer Science" OR "Construction & Building Technology" OR "Criminology & Penology" OR "Critical Care Medicine" OR "Crystallography" OR "Cultural Studies" OR "Dance" OR "Demography" OR "Dentistry, Oral Surgery & Medicine" OR "Dermatology" OR "Development Studies" OR "Education & Educational Research" OR "Electrochemistry" OR "Emergency Medicine" OR "Ethnic Studies" OR "Family Studies" OR "Film, Radio & Television" OR "Gastroenterology & Hepatology" OR "General & Internal Medicine" OR "Geochemistry & Geophysics" OR "Geriatrics & Gerontology" OR "Government & Law" OR "Health Care Sciences & Services" OR "Hematology" OR "History" OR "History & Philosophy of Science" OR "Imaging Science & Photographic Technology" OR "Information Science & Library Science" OR "Instruments & Instrumentation" OR "Integrative & Complementary Medicine" OR "International Relations" OR "Legal Medicine" OR "Linguistics" OR "Literature" OR "Mathematical & Computational Biology" OR "Mathematical Methods In Social Sciences" OR "Mathematics" OR "Mechanics" OR "Medical Ethics" OR "Medical Informatics" OR "Medical Laboratory Technology" OR "Metallurgy & Metallurgical Engineering" OR "Microscopy" OR "Music" OR "Nuclear Science & Technology" OR "Nursing" OR "Obstetrics & Gynecology" OR "Oncology" OR "Operations Research & Management Science" OR "Ophthalmology" OR "Optics" OR "Orthopedics" OR | *specify WoS research areas which will be excluded* |

"Otorhinolaryngology" OR "Pathology" OR "Pediatrics" OR "Philosophy" OR "Physics" OR "Polymer Science" OR "Public Administration" OR "Rehabilitation" OR "Religion" OR "Research & Experimental Medicine" OR "Respiratory System" OR "Rheumatology" OR "Robotics" OR "Social Issues" OR "Social Sciences Other Topics" OR "Social Work" OR "Sociology" OR "Sport Sciences" OR "Substance Abuse" OR "Surgery" OR "Telecommunications" OR "Theater" OR "Thermodynamics" OR "Transplantation" OR "Transportation" OR "Urology & Nephrology" OR "Women's Studies" OR "Anatomy & Morphology" OR "Astronomy & Astrophysics" OR "Energy & Fuels" OR "Infectious Diseases" OR "Materials Science" OR "Mineralogy" OR "Mining & Mineral Processing" OR "Neurosciences & Neurology" OR "Paleontology" OR "Pharmacology & Pharmacy" OR "Psychiatry" OR "Psychology" OR "Radiology, Nuclear Medicine & Medical Imaging" OR "Remote Sensing" OR "Spectroscopy" OR "Tropical Medicine" OR "Urban Studies" OR "Virology" OR "Biochemistry & Molecular Biology" OR "Cell Biology" OR "Endocrinology & Metabolism" OR "Genetics & Heredity" OR "Physiology")

29

Data figures

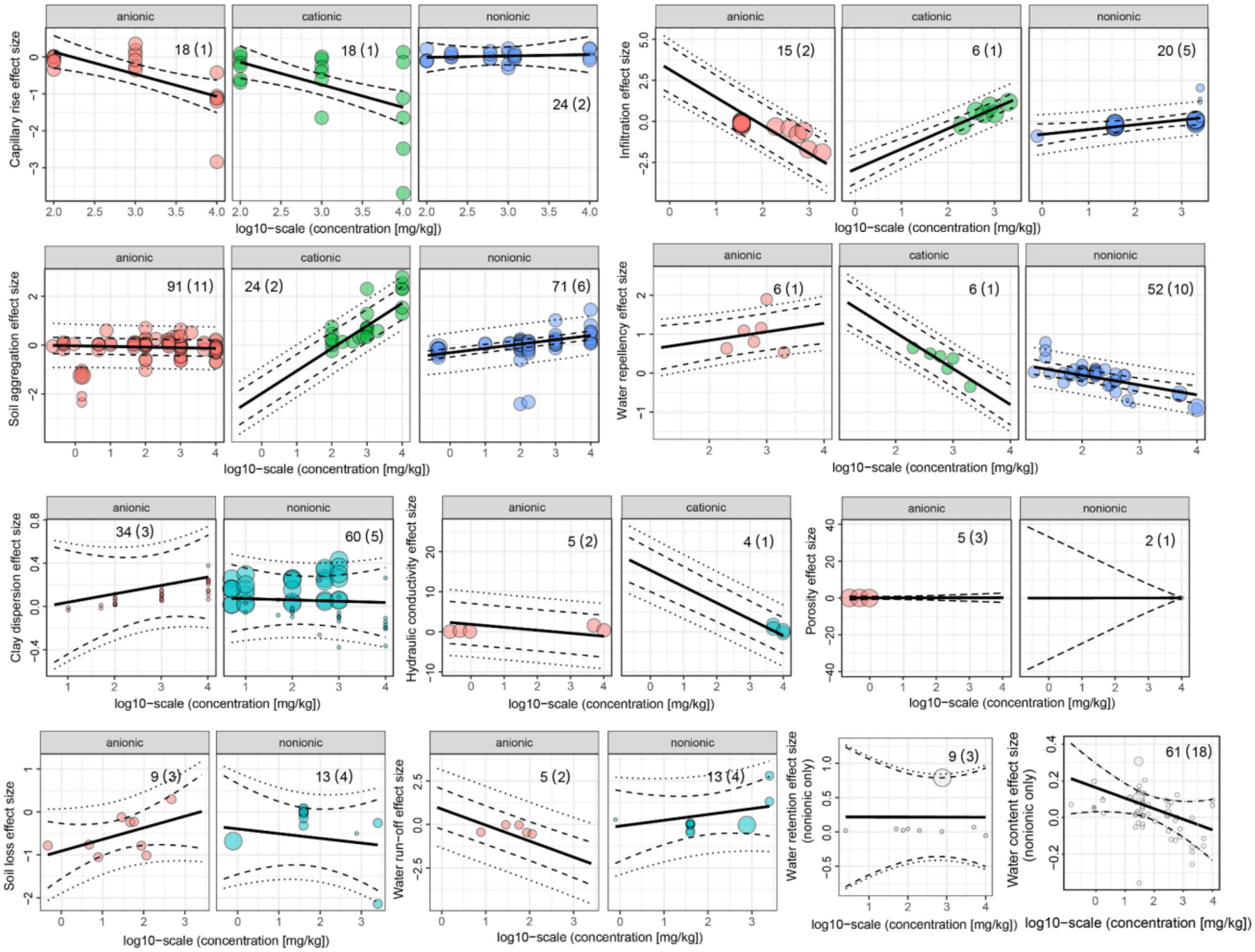

**Fig.S2.** Effect of surfactant concentration (log10-transformed in mg/kg) on individual soil hydrological and physical property effect sizes (soil aggregation, soil porosity, clay dispersion, soil water repellency (hydrophobicity), water run-off, soil loss (splash and detachment erosion), water infiltration, capillary rise, hydraulic conductivity, soil water retention and soil water content).The effect size estimate (solid line), the 95% confidence (dashed line) and prediction interval (dotted line) with individual effect sizes estimates in the background are shown. The number of effect sizes (k) are given together with the numbers of articles (number in brackets) from which these k effect sizes originate. Model outcomes are given in Table S4.

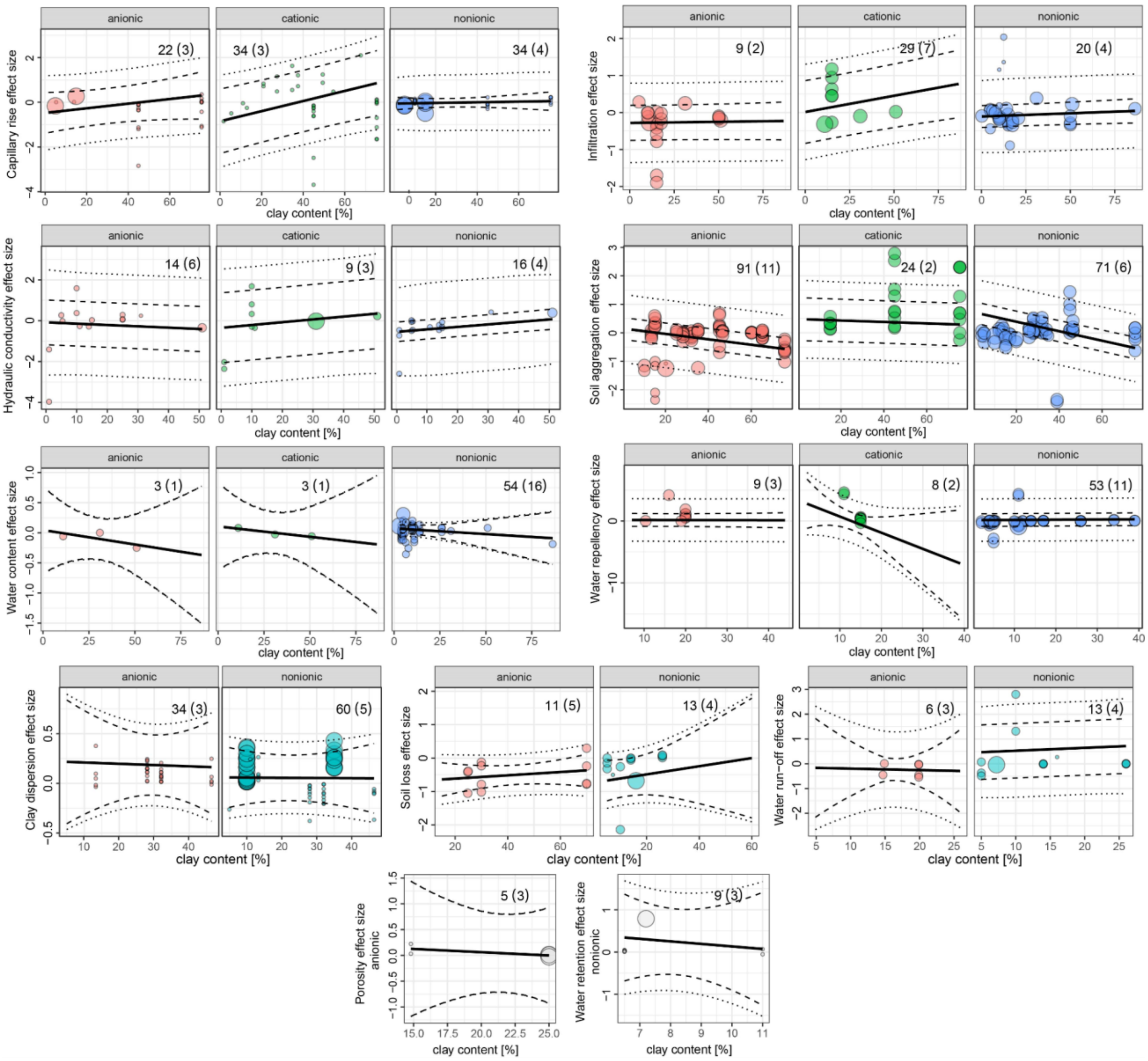

**Fig.S3.** Effect of clay content (in %) on individual soil hydrological and physical property effect sizes (soil aggregation, soil porosity, clay dispersion, soil water repellency (hydrophobicity), water run-off, soil loss (splash and detachment erosion), water infiltration, capillary rise, hydraulic conductivity, soil water retention and soil water content).The effect size estimate (solid line), the 95% confidence (dashed line) and prediction interval (dotted line) with individual effect sizes estimates in the background are shown. The number of effect sizes (k) are given together with the numbers of articles (number in brackets) from which these k effect sizes originate. Model outcomes are given in Table S5.

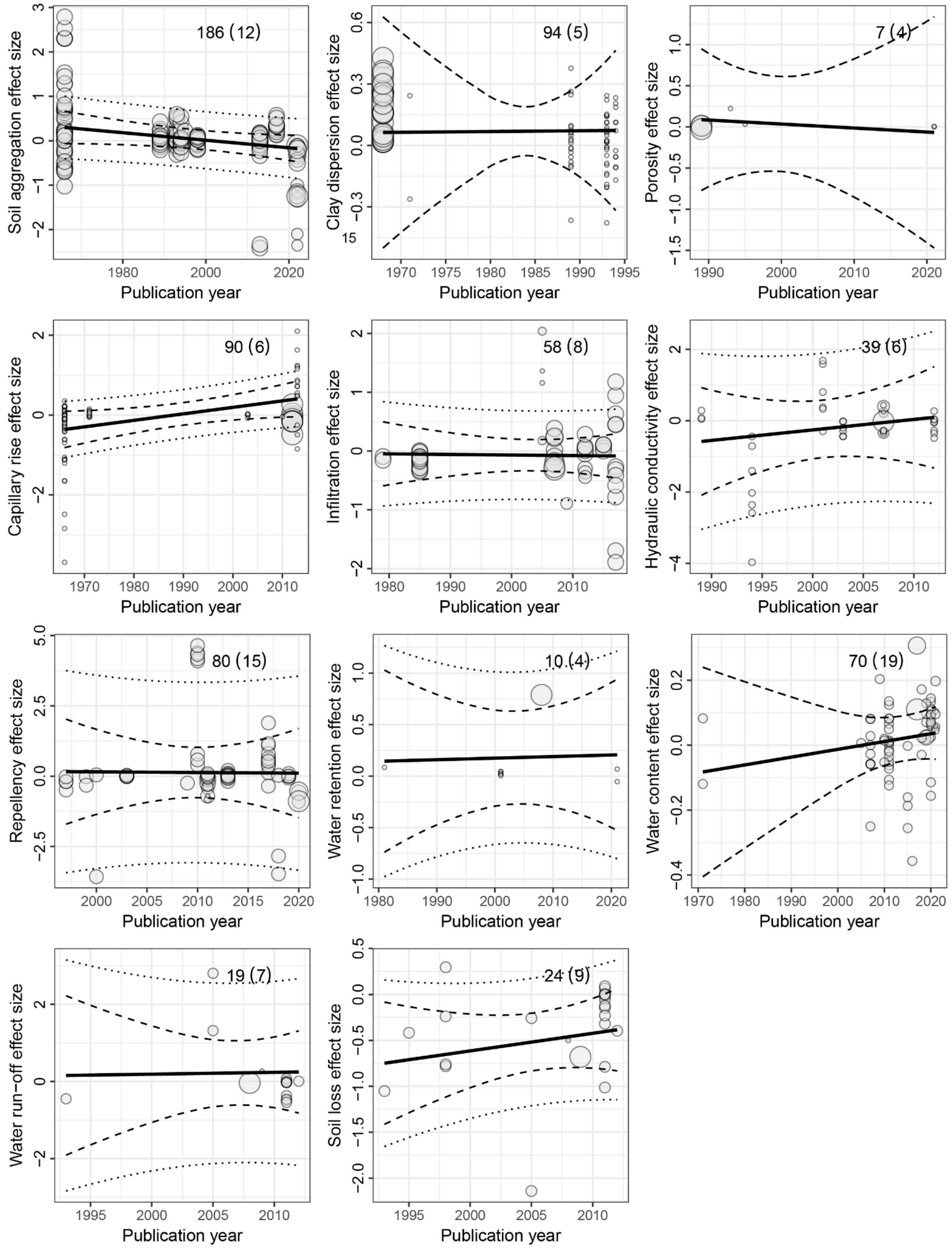

**Fig. S4.** Time-lag bias test for individual soil hydrological and physical property effect sizes (soil aggregation, soil porosity, clay dispersion, soil water repellency (hydrophobicity), water run-off, soil loss (splash and detachment erosion), water infiltration, capillary rise, hydraulic conductivity, soil water retention and soil water content).The effect size estimate (solid line), the 95% confidence (dashed line)

and prediction interval (dotted line) with individual effect sizes estimates in the background are shown. The number of effect sizes (k) are given together with the numbers of articles (number in brackets) from which these k effect sizes originate. Model outcomes are given in Table S6.

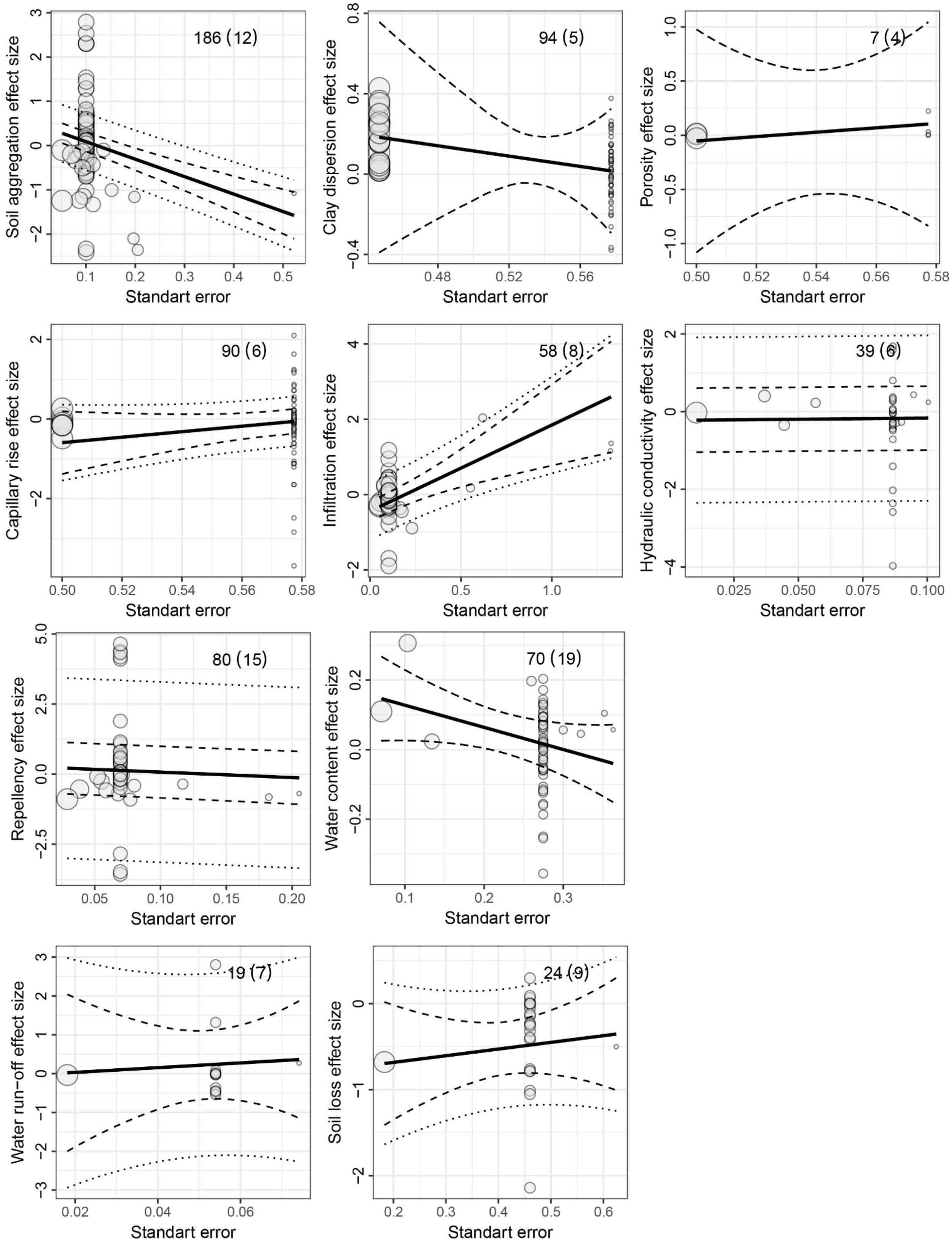

**Fig.S5.** Small study bias for individual soil hydrological and physical property effect sizes (soil aggregation, soil porosity, clay dispersion, soil water repellency (hydrophobicity), water run-off, soil loss (splash and detachment erosion), water infiltration, capillary rise, hydraulic conductivity, soil water retention and soil water content).The effect size estimate (solid line), the 95% confidence (dashed line) and prediction interval (dotted line) with individual effect sizes estimates in the background are shown. The number of effect sizes (k) are given together with the numbers of articles (number in brackets) from which these k effect sizes originate. Model outcomes are given in Table S6. (There is no plot for water retention because there was only one type of SE available: “0.104169”.)

Data tables

Table S1. Overall effect of surfactant application on individual soil hydrological and physical property and process effect sizes (soil aggregation, soil porosity, clay dispersion, soil water repellency (hydrophobicity), water run-off, soil loss (splash and detachment erosion), water infiltration, capillary rise, hydraulic conductivity, soil water retention and soil water content).

| effect size | estimate | lowerCL | upperCL | lowerPR | upperPR | pval | number of effect sizes | number of articles |
|---|---|---|---|---|---|---|---|---|
| soil aggregation | 0.04287466 | -0.2011479 | 0.2868972 | -0.6855701 | 0.7713194 | 0.7293 | 186 | 12 |
| clay dispersion | 0.09070305 | -0.05647471 | 0.2378808 | -0.1162069 | 0.297613 | 0.2241 | 94 | 5 |
| porosity | 0.03228551 | -0.4671883 | 0.5317593 | -0.4671883 | 0.5317593 | 0.8795 | 7 | 4 |
| capillary rise | 0.02158198 | -0.340967 | 0.384131 | -0.8321847 | 0.8753486 | 0.9061 | 90 | 6 |
| infiltration | -0.1069806 | -0.2739266 | 0.05996547 | -0.5628356 | 0.3488745 | 0.2046 | 58 | 8 |
| hydraulic conductivity | -0.2328647 | -0.9756506 | 0.5099212 | -2.196857 | 1.731128 | 0.5295 | 39 | 6 |
| water repellency | 0.1298417 | -0.7033987 | 0.9630822 | -2.898882 | 3.158566 | 0.7573 | 80 | 15 |
| water retention | 0.1439406 | -0.1709493 | 0.4588304 | -0.3354991 | 0.6233802 | 0.3281 | 10 | 4 |
| water content | 0.04412739 | -0.01876772 | 0.1070225 | -0.05976307 | 0.1480178 | 0.1661 | 70 | 19 |
| water run-off | 0.214417 | -0.4565844 | 0.8854184 | -1.681112 | 2.109946 | 0.5105 | 19 | 7 |
| soil loss | -0.5263052 | -0.8083731 | -0.2442373 | -1.153666 | 0.1010558 | **0.0008** | 24 | 9 |

estimate = effect size point estimate
lowerCL = lower 95% confidence interval
upperCL = upper 95% confidence interval
lowerPR = lower 95% prediction interval
upperPR = upper 95% prediction interval
pval = p-value

Table S2. Effect of 'setting' on surfactant-mediated effects on individual soil hydrological and physical property and process effect sizes (soil aggregation, soil porosity, clay dispersion, soil water repellency (hydrophobicity), water run-off, soil loss (splash and detachment erosion), water infiltration, capillary rise, hydraulic conductivity, soil water retention and soil water content).

| effect size | level | estimate | lowerCL | upperCL | lowerPR | upperPR | test of moderators | number of effect sizes | number of articles |
|---|---|---|---|---|---|---|---|---|---|
| soil aggregation | Field | -0.2127566 | -0.49167535 | 0.06616207 | -0.8935178 | 0.4680045 | **p-val = 0.0035** | 22 | 3 |
| | Lab | 0.1566626 | -0.07710492 | 0.39043007 | -0.5068781 | 0.8202032 | | 164 | 9 |
| clay dispersion | Field | -0.00970904 | -0.83845347 | 0.8190354 | -0.8560018 | 0.8365838 | p-val = 0.8069 | 2 | 1 |
| | Lab | 0.09440554 | -0.06223157 | 0.2510426 | -0.129168 | 0.3179791 | | 92 | 4 |
| porosity | Field only | NA | NA | NA | NA | NA | NA | 7 | 4 |
| capillary rise | | | | | | | | | |
| | Lab only | NA | NA | NA | NA | NA | NA | 90 | 6 |
| infiltration | Field | 0.05251146 | -1.8852395 | 1.99026237 | -3.2454227 | 3.35044562 | p-val = 0.8783 | 5 | 2 |
| | Lab | -0.09637027 | -0.1745312 | -0.01820931 | -0.2861108 | 0.09337022 | | 53 | 6 |
| hydraulic conductivity | Field | 0.1347238 | -0.9038 | 1.1732475 | -1.33047 | 1.599918 | p-val = 0.5180 | 3 | 1 |
| | Lab | -0.3062437 | -1.198154 | 0.5856665 | -2.490104 | 1.877616 | | 36 | 5 |
| water repellency | Field | -0.2653849 | -0.7346131 | 0.2038433 | -1.669733 | 1.138963 | p-val = 0.4602 | 26 | 8 |
| | Lab | 0.3152757 | -1.1699586 | 1.8005101 | -3.88514 | 4.515691 | | 54 | 7 |
| water retention | Field | 0.00716077 | -0.7183017 | 0.7326232 | -0.8735345 | 0.887856 | p-val = 0.5613 | 2 | 1 |
| | Lab | 0.24232463 | -0.2816319 | 0.7662812 | -0.6309175 | 1.115567 | | 8 | 3 |

| | | | | | | | | | |
|---|---|---|---|---|---|---|---|---|---|
| water content | Field | 0.07156995 | -0.00514264 | 0.14828254 | -0.03721723 | 0.18035714 | p-val = 0.2188 | 38 | 14 |
| | Lab | -0.00615936 | -0.10481384 | 0.09249513 | -0.10481384 | 0.09249513 | | 32 | 5 |
| water run-off | Field | 0.4732396 | -0.6880097 | 1.63448895 | -2.1206533 | 3.0671326 | p-val = 0.2930 | 5 | 4 |
| | Lab | -0.1286516 | -0.2723866 | 0.01508329 | -0.4125557 | 0.1552524 | | 14 | 3 |
| soil loss | Field | -0.7559531 | -1.0394467 | -0.47245953 | -1.039447 | -0.4724595 | **p-val = 0.0306** | 6 | 5 |
| | Lab | -0.2871333 | -0.5980819 | 0.02381537 | -0.767294 | 0.1930275 | | 18 | 4 |

estimate = effect size point estimate
level = moderator level
lowerCL = lower 95% confidence interval
upperCL = upper 95% confidence interval
lowerPR = lower 95% prediction interval
upperPR = upper 95% prediction interval
pval = p-value

Table S3. Effect of 'surfactant charge' on surfactant-mediated effects on individual soil hydrological and physical property and process effect sizes (soil aggregation, soil porosity, clay dispersion, soil water repellency (hydrophobicity), water run-off, soil loss (splash and detachment erosion), water infiltration, capillary rise, hydraulic conductivity, soil water retention and soil water content).

| **effect size** | **level** | **estimate** | **lowerCL** | **upperCL** | **lowerPR** | **upperPR** | **test of moderators** | **number of effect sizes** | **number of articles** |
|---|---|---|---|---|---|---|---|---|---|
| soil aggregation | Anionic | -0.1188487 | -0.4048715 | 0.1671741 | -1.0420887 | 0.8043912 | p-val = 0.1009 | 91 | 11 |
| | Cationic | 0.6008489 | -0.2667773 | 1.468475 | -0.9479215 | 2.1496193 | | 24 | 2 |
| | Nonionic | 0.1146177 | -0.1186391 | 0.3478745 | -0.4601276 | 0.6893629 | | 71 | 6 |
| clay dispersion | Anionic | 0.18359925 | -0.1091185 | 0.476317 | -0.2140476 | 0.5812461 | p-val = 0.3578 | 34 | 3 |
| | Cationic | NA | NA | NA | NA | NA | | | |
| | Nonionic | 0.05620576 | -0.1681118 | 0.2805233 | -0.2941621 | 0.4065736 | | 60 | 5 |
| porosity | Anionic | 0.04200242 | -0.5638895 | 0.6478944 | -0.5638895 | 0.6478944 | p-val = 0.9376 | 5 | 3 |
| | Cationic | NA | NA | NA | NA | NA | | 2 | 1 |
| | Nonionic | 0.00313481 | -1.0494444 | 1.055714 | -1.0525787 | 1.0588483 | | | |
| capillary rise | Anionic | -0.26229921 | -0.6575303 | 0.1329319 | -0.8525607 | 0.3279622 | p-val = 0.5533 | 22 | 3 |
| | Cationic | -0.03505712 | -0.9809498 | 0.9108356 | -1.8217707 | 1.7516565 | | 34 | 3 |
| | Nonionic | -0.0215939 | -0.2128516 | 0.1696638 | -0.2128516 | 0.1696638 | | 34 | 4 |
| infiltration | Anionic | -0.27009044 | -0.7402264 | 0.2000456 | -1.3169648 | 0.776784 | p-val = 0.5888 | 20 | 4 |
| | Cationic | 0.17807814 | -0.5965847 | 0.9527409 | -1.1611622 | 1.5173185 | | 9 | 2 |
| | Nonionic | -0.07609239 | -0.3698206 | 0.2176358 | -0.8581325 | 0.7059477 | | 29 | 7 |

| | | | | | | | | | |
|---|---|---|---|---|---|---|---|---|---|
| hydraulic conductivity | Anionic | -0.2339875 | -1.321374 | 0.8533993 | -3.076138 | 2.608163 | p-val = 0.9819 | 14 | 6 |
| | Cationic | -0.3339934 | -2.089655 | 1.4216685 | -3.825064 | 3.157077 | | 9 | 3 |
| | Nonionic | -0.3405252 | -0.963728 | 0.2826777 | -1.736785 | 1.055735 | | 16 | 4 |
| water repellency | Anionic | 0.6526025 | -0.458186 | 1.763391 | -2.303182 | 3.608387 | p-val = 0.5421 | 9 | 3 |
| | Cationic | 0.4413687 | -1.080426 | 1.963163 | -3.020158 | 3.902895 | | 8 | 2 |
| | Nonionic | 0.148199 | -0.7952044 | 1.091602 | -3.150754 | 3.447152 | | 63 | 14 |
| water retention | Anionic | 0.08463198 | -0.8036946 | 0.9729585 | -0.9274517 | 1.096716 | p-val = 0.7984 | 1 | 1 |
| | Cationic | NA | NA | NA | NA | NA | | | |
| | Nonionic | 0.20084984 | -0.2897194 | 0.6914191 | -0.6360501 | 1.03775 | | 9 | 3 |
| water content | Anionic | -0.1009512 | -0.43225788 | 0.2303555 | -0.4468731 | 0.2449707 | p-val = 0.6495 | 3 | 1 |
| | Cationic | 0.00091542 | -0.33039104 | 0.3322219 | -0.345006 | 0.3468369 | | 3 | 1 |
| | Nonionic | 0.05156992 | -0.01265463 | 0.1157945 | -0.0486227 | 0.1517625 | | 64 | 19 |
| water run-off | Anionic | -0.2352516 | -0.5107091 | 0.04020581 | -0.7773519 | 0.3068486 | p-val = 0.1533 | 6 | 3 |
| | Cationic | NA | NA | NA | NA | NA | | | |
| | Nonionic | 0.5512677 | -0.5241051 | 1.62664056 | -1.8515817 | 2.9541172 | | 13 | 4 |
| soil loss | Anionic | -0.508707 | -0.8112139 | -0.20620011 | -0.8112139 | -0.2062001 | p-val = 0.8350 | 11 | 5 |
| | Cationic | NA | NA | NA | NA | NA | | | |
| | Nonionic | -0.5736645 | -1.1384323 | -0.00889668 | -1.6680964 | 0.5207675 | | 13 | 4 |

estimate = effect size point estimate
level = moderator level

lowerCL = lower 95% confidence interval
upperCL = upper 95% confidence interval
lowerPR = lower 95% prediction interval
upperPR = upper 95% prediction interval
pval = p-value

Table S4. Effect of surfactant concentration (log10-transformed in mg/kg) on surfactant-mediated effects on individual soil hydrological and physical property and process effect sizes (soil aggregation, soil porosity, clay dispersion, soil water repellency (hydrophobicity), water run-off, soil loss (splash and detachment erosion), water infiltration, capillary rise, hydraulic conductivity, soil water retention and soil water content). Effects depicted for surfactant charges. See plots in Fig. S2.

| **effect size** | **surfactant charge** | | **estimate** | **lowerCL** | **upperCL** | **pval** | **number of effect sizes** | **number of articles** |
|---|---|---|---|---|---|---|---|---|
| soil aggregation | anionic | intercept | -0.0595 | -0.497 | 0.378 | 0.7876 | 91 | 11 |
| | | surfactant_concentration (log10[mg/kg]) | -0.0322 | -0.0545 | -0.01 | **0.0049** | | |
| | cationic | intercept | -1.9692 | -2.6684 | -1.2699 | <.0001 | 24 | 2 |
| | | surfactant_concentration (log10[mg/kg]) | 0.9242 | 0.8656 | 0.9829 | **<.0001** | | |
| | nonionic | intercept | -0.3485 | -0.5458 | -0.1512 | 0.0008 | 71 | 6 |
| | | surfactant_concentration (log10[mg/kg]) | 0.1805 | 0.1465 | 0.2145 | **<.0001** | | |
| clay dispersion | anionic | intercept | -0.1167 | -0.751 | 0.5176 | 0.7103 | 34 | 3 |
| | | surfactant_concentration (log10[mg/kg]) | 0.0777 | -0.1367 | 0.292 | 0.4659 | | |
| | cationic | intercept | NA | NA | NA | NA | | |
| | | surfactant_concentration (log10[mg/kg]) | NA | NA | NA | NA | | |
| | nonionic | intercept | 0.0838 | -0.3012 | 0.4689 | 0.6646 | 60 | 5 |
| | | surfactant_concentration (log10[mg/kg]) | -0.0119 | -0.1482 | 0.1244 | 0.8623 | | |
| porosity | anionic | intercept | 0.0336 | -0.7547 | 0.8219 | 0.9007 | 5 | 3 |
| | | surfactant_concentration (log10[mg/kg]) | 0.0229 | -0.6373 | 0.6832 | 0.919 | | |
| | cationic | intercept | NA | NA | NA | NA | | |
| | | surfactant_concentration (log10[mg/kg]) | NA | NA | NA | NA | | |

| | | | | | | | | |
|---|---|---|---|---|---|---|---|---|
| | nonionic | intercept | -0.077 | -132.8455 | 132.6914 | 0.9953 | 2 | 1 |
| | | surfactant_concentration (log10[mg/kg]) | 0.0208 | -34.4428 | 34.4844 | 0.9951 | | |
| capillary rise | anionic | intercept | 1.3355 | 0.2349 | 2.4361 | 0.0205 | 18 | 1 |
| | | surfactant_concentration (log10[mg/kg]) | -0.6015 | -0.9548 | -0.2482 | **0.0024** | | |
| | cationic | intercept | 1.0795 | -0.2928 | 2.4518 | 0.1148 | 18 | 1 |
| | | surfactant_concentration (log10[mg/kg]) | -0.6113 | -0.9647 | -0.258 | **0.0021** | | |
| | nonionic | intercept | -0.087 | -1.2131 | 1.039 | 0.8741 | 24 | 2 |
| | | surfactant_concentration (log10[mg/kg]) | 0.0371 | -0.3473 | 0.4214 | 0.8433 | | |
| infiltration | anionic | intercept | 3.1878 | 1.6575 | 4.7182 | 0.0006 | 15 | 2 |
| | | surfactant_concentration (log10[mg/kg]) | -1.6971 | -1.9849 | -1.4093 | **<.0001** | | |
| | cationic | intercept | -2.9117 | -4.0364 | -1.7869 | 0.002 | 6 | 1 |
| | | surfactant_concentration (log10[mg/kg]) | 1.2353 | 0.8623 | 1.6083 | **0.0008** | | |
| | nonionic | intercept | -0.7873 | -1.4496 | -0.1249 | 0.0224 | 20 | 5 |
| | | surfactant_concentration (log10[mg/kg]) | 0.2922 | 0.0434 | 0.541 | **0.0239** | | |
| hydraulic conductivity | anionic | intercept | 1.8526 | -4.5929 | 8.2982 | 0.4278 | 5 | 2 |
| | | surfactant_concentration (log10[mg/kg]) | -0.737 | -1.306 | -0.168 | **0.0259** | | |
| | cationic | intercept | 16.2963 | 11.406 | 21.1866 | 0.0048 | 4 | 1 |
| | | surfactant_concentration (log10[mg/kg]) | -4.0695 | -5.3078 | -2.8313 | **0.005** | | |
| | nonionic | intercept | NA | NA | NA | NA | | |

| | | | | | | | | |
|---|---|---|---|---|---|---|---|---|
| | | surfactant_concentration (log10[mg/kg]) | NA | NA | NA | NA | | |
| water repellency | anionic | intercept | 0.4022 | -0.649 | 1.4534 | 0.348 | 6 | 1 |
| | | surfactant_concentration (log10[mg/kg]) | 0.219 | -0.0323 | 0.4703 | 0.0728 | | |
| | cationic | intercept | 2.885 | 2.135 | 3.6351 | 0.0004 | 6 | 1 |
| | | surfactant_concentration (log10[mg/kg]) | -0.9238 | -1.1751 | -0.6725 | **0.0005** | | |
| | nonionic | intercept | 0.4488 | 0.1813 | 0.7163 | 0.0015 | 52 | 10 |
| | | surfactant_concentration (log10[mg/kg]) | -0.2519 | -0.3468 | -0.157 | **<.0001** | | |
| water retention | anionic | intercept | NA | NA | NA | NA | | |
| | | surfactant_concentration (log10[mg/kg]) | NA | NA | NA | NA | | |
| | cationic | intercept | NA | NA | NA | NA | | |
| | | surfactant_concentration (log10[mg/kg]) | NA | NA | NA | NA | | |
| | nonionic | intercept | 0.2189 | -0.9103 | 1.3481 | 0.6606 | 9 | 3 |
| | | surfactant_concentration (log10[mg/kg]) | -0.0014 | -0.3514 | 0.3485 | 0.9927 | | |
| water content | anionic | intercept | NA | NA | NA | NA | | |
| | | surfactant_concentration (log10[mg/kg]) | NA | NA | NA | NA | | |
| | cationic | intercept | NA | NA | NA | NA | | |
| | | surfactant_concentration (log10[mg/kg]) | NA | NA | NA | NA | | |
| | nonionic | intercept | 0.1632 | 0.0285 | 0.298 | 0.0185 | 61 | 18 |
| | | surfactant_concentration (log10[mg/kg]) | -0.0577 | -0.125 | 0.0097 | 0.0918 | | |
| water run-off | anionic | intercept | 0.8807 | -0.8357 | 2.597 | 0.201 | 5 | 2 |

| | | | | | | | | |
|---|---|---|---|---|---|---|---|---|
| | | surfactant_concentration (log10[mg/kg]) | -0.9203 | -1.2979 | -0.5427 | **0.0045** | | |
| | cationic | intercept | NA | NA | NA | NA | | |
| | | surfactant_concentration (log10[mg/kg]) | NA | NA | NA | NA | | |
| | nonionic | intercept | -0.0976 | -2.1689 | 1.9737 | 0.9193 | 13 | 4 |
| | | surfactant_concentration (log10[mg/kg]) | 0.3334 | -0.5407 | 1.2075 | 0.4191 | | |
| soil loss | anionic | intercept | -0.9082 | -1.6335 | -0.1829 | 0.0211 | 9 | 3 |
| | | surfactant_concentration (log10[mg/kg]) | 0.2703 | -0.1659 | 0.7066 | 0.1863 | | |
| | cationic | intercept | NA | NA | NA | NA | | |
| | | surfactant_concentration (log10[mg/kg]) | NA | NA | NA | NA | | |
| | nonionic | intercept | -0.3956 | -1.5079 | 0.7168 | 0.4503 | 13 | 4 |
| | | surfactant_concentration (log10[mg/kg]) | -0.1099 | -0.6154 | 0.3955 | 0.6415 | | |

estimate = effect size point estimate
level = moderator level
lowerCL = lower 95% confidence interval
upperCL = upper 95% confidence interval
pval = p-value

Table S5. Effect of clay content (%) on surfactant-mediated effects on individual soil hydrological and physical property and process effect sizes (soil aggregation, soil porosity, clay dispersion, soil water repellency (hydrophobicity), water run-off, soil loss (splash and detachment erosion), water infiltration, capillary rise, hydraulic conductivity, soil water retention and soil water content). Effects depicted for surfactant charges. See plots in Fig. S3.

| **effect size** | **surfactant charge** | | **estimate** | **lowerCL** | **upperCL** | **pval** | **number of effect sizes** | **number of articles** |
|---|---|---|---|---|---|---|---|---|
| soil aggregation | anionic | intercept | 0.1817 | -0.2579 | 0.6214 | 0.4137 | 91 | 11 |
| | | clay content [%] | -0.0093 | -0.0113 | -0.0074 | **<.0001** | | |
| | cationic | intercept | 0.8147 | -0.1729 | 1.8024 | 0.1012 | 24 | 2 |
| | | clay content [%] | -0.0026 | -0.0058 | 0.0006 | 0.1094 | | |
| | nonionic | intercept | 0.5054 | -0.0709 | 1.0818 | 0.0847 | 71 | 6 |
| | | clay content [%] | -0.0167 | -0.0188 | -0.0146 | **<.0001** | | |
| clay dispersion | anionic | intercept | 0.1396 | -0.5521 | 0.8313 | 0.6837 | 34 | 3 |
| | | clay content [%] | -0.0013 | -0.0234 | 0.0208 | 0.9065 | | |
| | cationic | intercept | NA | NA | NA | NA | | |
| | | clay content [%] | NA | NA | NA | NA | | |
| | nonionic | intercept | 0.0601 | -0.2789 | 0.3991 | 0.7239 | 60 | 5 |
| | | clay content [%] | -0.0002 | -0.0112 | 0.0108 | 0.971 | | |
| porosity | anionic | intercept | 0.3141 | -3.1499 | 3.7782 | 0.7917 | 5 | 3 |
| | | clay content [%] | -0.0126 | -0.1692 | 0.144 | 0.8145 | | |
| | cationic | intercept | NA | NA | NA | NA | | |
| | | clay content [%] | NA | NA | NA | NA | | |
| | nonionic | intercept | NA | NA | NA | NA | | |
| | | clay content [%] | NA | NA | NA | NA | | |
| capillary rise | anionic | intercept | -0.477 | -1.4334 | 0.4794 | 0.3106 | 22 | 3 |

| | | | | | | | | |
|---|---|---|---|---|---|---|---|---|
| | | clay content [%] | 0.0104 | -0.0058 | 0.0265 | 0.196 | | |
| | cationic | intercept | -0.8497 | -2.3281 | 0.6287 | 0.2504 | 34 | 3 |
| | | clay content [%] | 0.0226 | 0.0103 | 0.0349 | **0.0007** | | |
| | nonionic | intercept | -0.0601 | -0.3444 | 0.2242 | 0.6697 | 34 | 4 |
| | | clay content [%] | 0.0014 | -0.0063 | 0.0092 | 0.7059 | | |
| infiltration | anionic | intercept | -0.2816 | -0.7776 | 0.2144 | 0.2485 | 20 | 4 |
| | | clay content [%] | 0.0006 | -0.0025 | 0.0037 | 0.6887 | | |
| | cationic | intercept | 0.013 | -0.9867 | 1.0127 | 0.9763 | 9 | 2 |
| | | clay content [%] | 0.0088 | 0.0027 | 0.0149 | **0.0113** | | |
| | nonionic | intercept | -0.1097 | -0.4123 | 0.1929 | 0.4635 | 29 | 7 |
| | | clay content [%] | 0.0018 | -0.0004 | 0.004 | 0.1134 | | |
| hydraulic conductivity | anionic | intercept | -0.201 | -1.3418 | 0.9398 | 0.7078 | 14 | 6 |
| | | clay content [%] | -0.0064 | -0.0115 | -0.0013 | **0.0176** | | |
| | cationic | intercept | -0.7214 | -2.5927 | 1.1499 | 0.3923 | 9 | 3 |
| | | clay content [%] | 0.0138 | 0.0082 | 0.0194 | **0.0007** | | |
| | nonionic | intercept | -0.5437 | -1.066 | -0.0213 | 0.0424 | 16 | 4 |
| | | clay content [%] | 0.0121 | 0.0076 | 0.0166 | **<.0001** | | |
| water repellency | anionic | intercept | 1.7485 | -1.2403 | 4.7374 | 0.2091 | 9 | 3 |
| | | clay content [%] | -0.0016 | -0.0286 | 0.0254 | 0.8949 | | |
| | cationic | intercept | 16.0366 | 13.1575 | 18.9156 | <.0001 | 8 | 2 |
| | | clay content [%] | -1.0501 | -1.2686 | -0.8316 | **<.0001** | | |
| | nonionic | intercept | 0.0417 | -1.0893 | 1.1728 | 0.9413 | 53 | 11 |
| | | clay content [%] | 0.0037 | 0.0012 | 0.0063 | **0.0053** | | |
| water retention | anionic | intercept | NA | NA | NA | NA | | |

| | | | | | | | | |
|---|---|---|---|---|---|---|---|---|
| | | clay content [%] | NA | NA | NA | NA | | |
| | cationic | intercept | NA | NA | NA | NA | | |
| | | clay content [%] | NA | NA | NA | NA | | |
| | nonionic | intercept | 0.7322 | -2.5472 | 4.0115 | 0.6139 | 9 | 3 |
| | | clay content [%] | -0.06 | -0.448 | 0.328 | 0.7256 | | |
| water content | anionic | intercept | 0.0456 | -4.2962 | 4.3875 | 0.9155 | 3 | 1 |
| | | clay content [%] | -0.0048 | -0.1282 | 0.1186 | 0.7082 | | |
| | cationic | intercept | 0.1057 | -4.2361 | 4.4476 | 0.8089 | 3 | 1 |
| | | clay content [%] | -0.0034 | -0.1268 | 0.1199 | 0.783 | | |
| | nonionic | intercept | 0.0725 | -0.0151 | 0.16 | 0.1028 | 54 | 16 |
| | | clay content [%] | -0.0019 | -0.0074 | 0.0037 | 0.5002 | | |
| water run-off | anionic | intercept | -0.1354 | -3.7302 | 3.4594 | 0.9217 | 6 | 3 |
| | | clay content [%] | -0.006 | -0.22 | 0.208 | 0.9418 | | |
| | cationic | intercept | NA | NA | NA | NA | | |
| | | clay content [%] | NA | NA | NA | NA | | |
| | nonionic | intercept | 0.4102 | -0.7278 | 1.5482 | 0.4444 | 13 | 4 |
| | | clay content [%] | 0.0117 | 0.0071 | 0.0163 | **0.0002** | | |
| soil loss | anionic | intercept | -0.6635 | -1.2621 | -0.065 | 0.0334 | 11 | 5 |
| | | clay content [%] | 0.0049 | -0.0105 | 0.0203 | 0.4874 | | |
| | cationic | intercept | NA | NA | NA | NA | | |
| | | clay content [%] | NA | NA | NA | NA | | |
| | nonionic | intercept | -0.7323 | -1.5023 | 0.0377 | 0.0603 | 13 | 4 |
| | | clay content [%] | 0.0122 | -0.0261 | 0.0506 | 0.4978 | | |

estimate = effect size point estimate
level = moderator level
lowerCL = lower 95% confidence interval
upperCL = upper 95% confidence interval
pval = p-value

Table S6. Model outcome for time-lag (publication year) and small study (sample size, SE) bias analyses (see Fig.S4 and S5).

| effect size | | estimate | lowerCL | upperCL | pval | number of effect sizes | number of articles |
|---|---|---|---|---|---|---|---|
| soil aggregation | intercept | 17.4032 | -0.1829 | 34.9894 | 0.0524 | 186 | 12 |
| | year | -0.0085 | -0.0173 | 0.0003 | 0.0584 | | |
| | sei | -3.9435 | -5.0674 | -2.8196 | **<.0001** | | |
| clay dispersion | intercept | 0.0281 | -65.1472 | 65.2034 | 0.9993 | 94 | 5 |
| | year | 0.0004 | -0.0341 | 0.0349 | 0.9831 | | |
| | sei | -1.2865 | -7.6734 | 5.1004 | 0.69 | | |
| porosity | intercept | 8.6585 | -102.4661 | 119.7832 | 0.8393 | 7 | 4 |
| | year | -0.0049 | -0.0642 | 0.0544 | 0.8311 | | |
| | sei | 2.019 | -18.3743 | 22.4122 | 0.797 | | |
| capillary rise | intercept | -36.2862 | -68.5842 | -3.9882 | 0.0281 | 90 | 6 |
| | year | 0.0163 | 0.0016 | 0.031 | **0.0304** | | |
| | sei | 6.8949 | -3.3976 | 17.1874 | 0.1865 | | |
| infiltration | intercept | 1.4803 | -37.1781 | 40.1386 | 0.9391 | 58 | 8 |
| | year | -0.001 | -0.0203 | 0.0183 | 0.921 | | |
| | sei_inf | 2.288 | 1.0589 | 3.517 | **0.0005** | | |
| hydraulic conductivity | intercept | -59.1816 | -268.1073 | 149.7442 | 0.5692 | 39 | 6 |
| | year | 0.0294 | -0.075 | 0.1338 | 0.571 | | |
| | sei | 0.5858 | -0.3393 | 1.5109 | 0.2073 | | |
| water repellency | intercept | 4.8613 | -248.057 | 257.7797 | 0.9696 | 80 | 15 |
| | year | -0.0023 | -0.1281 | 0.1236 | 0.9713 | | |
| | sei_repel | -1.9492 | -3.4987 | -0.3997 | **0.0144** | | |

| | | | | | | | |
|---|---|---|---|---|---|---|---|
| water retention | intercept | -2.8684 | -68.9695 | 63.2328 | 0.9228 | 10 | 4 |
| | year | 0.0015 | -0.0315 | 0.0345 | 0.9179 | | |
| | sei | NA | NA | NA | NA | | |
| water content | intercept | -4.6485 | -19.712 | 10.415 | 0.54 | 70 | 19 |
| | year | 0.0024 | -0.0051 | 0.0099 | 0.5224 | | |
| | sei | -0.6372 | -1.3128 | 0.0384 | 0.0641 | | |
| water run-off | intercept | -9.6209 | -279.4952 | 260.2535 | 0.9407 | 19 | 7 |
| | year | 0.0047 | -0.1297 | 0.1392 | 0.9413 | | |
| | sei | 6.1047 | -47.3371 | 59.5465 | 0.8117 | | |
| soil loss | intercept | -39.2646 | -133.916 | 55.3868 | 0.3981 | 24 | 9 |
| | year | 0.0191 | -0.0279 | 0.0662 | 0.4072 | | |
| | sei | 0.7764 | -1.9362 | 3.489 | 0.5581 | | |

estimate = effect size point estimate
level = moderator level
lowerCL = lower 95% confidence interval
upperCL = upper 95% confidence interval
pval = p-value
